# PREpiBind: Protein Representation-integrated Epitope–MHC Class II Binding Prediction

**DOI:** 10.64898/2026.09.15.751749

**Authors:** David Hyunyoo Jang, Dongwoo Kim, Untaek Hwang, Byungho Park, Yoonjoo Choi, Juyong Lee

## Abstract

**Motivation:** Accurate peptide–MHC class II (pMHC-II) binding prediction is complicated by MHC polymorphism and context-dependent peptide recognition. Protein representations are usually evaluated in different pipelines, obscuring their contribution to predictive performance.

**Results:** We present PREpiBind, a joint-attention framework comparing ten protein representations under a fixed architecture and identical data splits across qualitative binding, mass spectrometry, and thresholded IC50 datasets. PLMs generally achieved the strongest pooled performance; ESM3 Large reached a ROC-AUC of 0.927 *±* 0.002 on the qualitative dataset. PREpiBind PLM variants exceeded evaluated reference methods on qualitative and mass-spectrometry benchmarks, whereas NetMHCIIpan-4.3 was higher on thresholded IC50 using its binding-affinity head. PLM advantages narrowed in allele-wise and held-out MHC-molecule evaluations, and H2-out rankings depended on aggregation, indicating scenario-dependent representation performance.

**Availability and implementation:** Source code and the training and evaluation datasets are available at https://github.com/daylight-00/PREpiBind and archived at https://doi.org/10.5281/zenodo.22934281; trained model checkpoints are at https://huggingface.co/daylight-00/prepibind. Implemented in Python and released under the MIT License.

**Supplementary information:** Supplementary data are available online.

## Introduction

Major histocompatibility complex class II (MHC-II) molecules play key roles in adaptive immunity. They present processed peptides from external antigens to CD4^+^ T cells, initiating immune responses such as vaccine-induced protection, anti-tumor responses, and the regulation of autoimmunity [1–3]. MHC-II molecules are expressed on antigen-presenting cells and sample peptides from endocytosed proteins via tightly regulated antigen processing pathways. These pathways involve proteolytic cleavage, peptide editing, and selection by chaperones, ultimately ensuring that only appropriately processed peptides are displayed to T cells [1]. The functional importance of this process is highlighted in clinical contexts including vaccine development, neoantigen discovery for cancer immunotherapy, and the study of autoimmune disorders [2–4].

Despite significant progress in the field, achieving highly accurate and generalized prediction of peptide–MHC-II (pMHC-II) binding remains a complex challenge, particularly compared to MHC-I. MHC-II molecules possess an open-ended binding groove that accommodates peptides of varying lengths (typically 13–25 amino acids) and allows for alternative binding registers [1]. The binding cores of MHC-II molecules are often ambiguous, as these peptides can occupy multiple binding registers within the groove [5, 6], and flanking residues outside the 9-mer core modulate both binding affinity and T cell recognition [7, 8]. Furthermore, human leukocyte antigen (HLA) genes are highly polymorphic [9]. This diversity results in allele-specific differences in peptide preferences and population-level immune responses [9, 10]. Recent studies have also revealed alternative binding modes for certain allotypes, further complicating the prediction task [6, 11, 12]. Such biological and structural complexity underlying pMHC-II interaction poses a formidable barrier to in silico prediction.

While established computational models have driven significant progress in pMHC-II binding prediction, they retain notable limitations [12–15]. Many rely on static or simplified input encodings [13, 14], and models built on them can generalize poorly across diverse alleles or miss context-dependent effects, which has motivated architectures designed to capture the binding core together with its context [16, 17]. Moreover, the reliance on highly imbalanced training data, heavily clustered around a few common alleles, increases the risk of population bias in model predictions [18]. This creates a significant gap between in silico predictions and their practical application in diverse populations.

The field of computational immunology is transforming rapidly due to advances in protein language models (PLMs) and 3D structure prediction methods [19, 20]. PLMs such as ESMC and ESM3, trained on billions of protein sequences, produce evolutionary context-aware embeddings that encode deep evolutionary and structural information [19, 21, 22]. Concurrently, biomolecular 3D structure-prediction models such as AlphaFold 3 (AF3), Chai-1 and Boltz-1 generate residue-level and pairwise representations with rich inter-chain interaction information [20, 23–25]. Recent work has begun to incorporate these pretrained representations into pMHC-II prediction. pMHChat encodes MHC molecules with a fine-tuned MSA-specialized language model alongside ESM-2 [26] peptide features in a hypergraph interaction model [27]. AlphaFold-based methods have been shown to complement sequence-based tools in locating pMHC-II binding cores [28]. However, these studies evaluate representations within distinct task-specific modeling pipelines rather than comparing broad representation families under a common downstream architecture and identical data splits. Such a comparison, spanning substitution-matrix, structure-prediction-derived, and modern PLM representations, has to our knowledge not been reported for pMHC-II binding.

To address this gap, we developed PREpiBind (**<u>P</u>**rotein **<u>R</u>**epresentation-integrated **<u>Epi</u>**tope-MHC class II **<u>Bind</u>**ing prediction), a unified, modular, and open-source deep learning prediction method. PREpiBind was designed to isolate the contribution of protein representations by holding the downstream architecture, training protocol, and data partitions fixed. To compare the effect of protein representations for pMHC-II binding prediction, we used the PREpiBind architecture to evaluate ten embedding strategies, ranging from classical sequence-based features (BLOSUM62) [13] to modern transformer-based protein language models (ESMC, ESM3) [21, 22] and structure-prediction-derived embeddings (AlphaFold 3, Boltz-1, Chai-1) [23–25]. We assessed predictive performance across three rigorous, assay-derived settings: qualitative binding labels, mass spectrometry (MS)-derived ligandomics, and quantitative IC50 measurements, all drawn from IEDB [29]. We then benchmarked our models against widely-used tools, including NetMHCIIpan-4.3 [14], MixMHC2pred-2.0 [12], and DeepNeo [15]. In the evaluated benchmark settings, ESM3-based PREpiBind models yielded higher ROC-AUC values than the included baseline methods in several analyses. Finally, leave-one-molecule-out (LOMO) and species-out validation schemes were used to examine performance on withheld alleles and mouse H2 targets, addressing the allele-imbalance bias documented for MHC binding predictors [18].

## Materials and Methods

### Dataset Formulation

All pMHC binding data were retrieved from the Immune Epitope Database (IEDB) [29] as of April 21, 2025, restricted to linear, unmutated 15-mer peptides for three human MHC-II serotypes (HLA-DR, HLA-DQ, HLA-DP) and murine H2. 15-mers are the largest single-length cohort in the IEDB, contain the canonical 9-mer core together with its flanking residues [5, 8], and keep peptide length fixed across representation methods. Human MHC-II sequences were taken from the IPD-IMGT/HLA Database [30–32], release 3.59.0, obtained from its public distribution at https://github.com/ANHIG/IMGTHLA, and murine H2 sequences from UniProt [33]; both were truncated to the peptide-binding domains using InterPro domain annotations retrieved from UniProt [34]. Filtering criteria, allele-specific handling and domain coordinates are given in Supplementary Text 1.1 and Supplementary Table S4.

Three evaluation settings were assembled — qualitative binding, mass spectrometry (MS) ligandomics and quantitative IC50 — comprising 161,223, 111,444 and 47,154 pMHC entries (Table 1). Qualitative labels carry the primary analyses, because they supply experimentally confirmed non-binders alongside binders and pool several assay platforms. MS elution instead reports peptides actually presented, which reflects antigen processing and surface stabilization as well as binding, and yields no verified non-binders. We therefore augmented the IEDB MS positives with experimentally measured negatives from the Qualitative dataset rather than sampling proteome-derived decoys as the published tools do [12, 14]. Proteome-derived decoys can introduce label noise and physicochemical bias [35]. Our choice instead creates a dependency between the Qualitative and MS settings, which we return to in the Discussion. IC50 measurements were binarised at 500 nM and at 1,000 nM, following the thresholds established for MHC-II binding assays [36].

**Table 1.** Dataset composition by MHC type. Parentheses give each serotype’s share of the samples counted in that row. MS serotype counts refer to positive eluted ligands; **Total** and **Positive** refer to the full dataset used for training and evaluation. The two IC50 percentages are the positive fractions at *<*500 and *<*1,000 nM.

| Dataset | HLA-DP | HLA-DQ | HLA-DR | H2 | Total | Positive (%) |
| --- | --- | --- | --- | --- | --- | --- |
| Qualitative | 27,293 (16.9) | 19,160 (11.9) | 111,650 (69.3) | 3,120 (1.9) | 161,223 | 58.1 |
| MS | 20,667 (47.1) | 4,983 (11.4) | 16,435 (37.5) | 1,783 (4.1) | 111,444 | 39.4 |
| IC50 | 4,738 (10.0) | 5,607 (11.9) | 35,645 (75.6) | 1,164 (2.5) | 47,154 | 36.7 / 46.0 |

The three settings share peptides and negatives and are therefore not additive: the Qualitative dataset contains every MS positive by construction, together with 99.6% of the IC50 *<* 500 nM positives and 82.9% of its negatives (99.4% of positives at 1,000 nM). Serotype composition also differs between them (Table 1), HLA-DR dominating the Qualitative and IC50 datasets and HLA-DP the MS dataset, with murine H2 smallest throughout; the intersections are given in Supplementary Figure S1.

### Training and Evaluation Datasets

Each dataset was partitioned 70:30 into a training set and an independent test set by stratified sampling on class label and HLA serotype. Partition sizes are given in Supplementary Table S1, and class and MHC-type composition in Table 1 and Supplementary Text 1.1. IC50 was additionally stratified across affinity bins (*<*500 nM, 500–1,000 nM, *≥*1,000 nM). Each training set was then split into five cross-validation folds under the same criteria (Supplementary Table S1 and Supplementary Text 1.1). Cross-validation was confined to the training partition. The independent test set took no part in fitting, early stopping or checkpoint selection. Therefore the benchmark results reported for each dataset, including the allele-wise and serotype breakdowns, were computed on the independent test set and not as an average over the cross-validation folds. The LOMO and species-out analyses use their own held-out partitions. Preventing leakage matters here because one entry pairs a peptide with MHC-II *α*- and *β*-chains and some entries carry no explicit *α*-chain annotation. We therefore de-duplicated records differing only by the presence or absence of an *α*-chain, and assigned all entries sharing an identical peptide and *β*-chain sequence as a group to either training or test, so no pMHC binding context appears in both.

Sequence homology was controlled at two levels, because the prediction unit here is a peptide–MHC context and not a peptide alone. At the pair level, the group assignment above keeps identical pMHC contexts out of the test partition. At the peptide level, generalization was measured on the subset of test epitopes sharing no exact 9-mer with any training epitope, the 9-mer core being the recognition unit in MHC-II and the unit NetMHCIIpan partitions on [14]. Global identity clustering was evaluated and not adopted: it left substantial 9-mer overlap while sharply reducing the training set and distorting class balance (Supplementary Text 1.5).

### Model Development

#### Model Architecture

The overall architecture is shown in Figure 1. The MHC and epitope embeddings enter two separate encoder blocks of two transformer layers each, are concatenated along the sequence dimension, pass through a single self-attention interaction block, and are mean-pooled and fed to a two-layer prediction head trained with binary cross-entropy (BCEWithLogitsLoss). Sequences are dynamically padded within each batch and the padding is masked in every attention operation. Layer composition, regularization and parameter counts are given in Supplementary Text 1.2 and Supplementary Table S6.

**Fig. 1.**
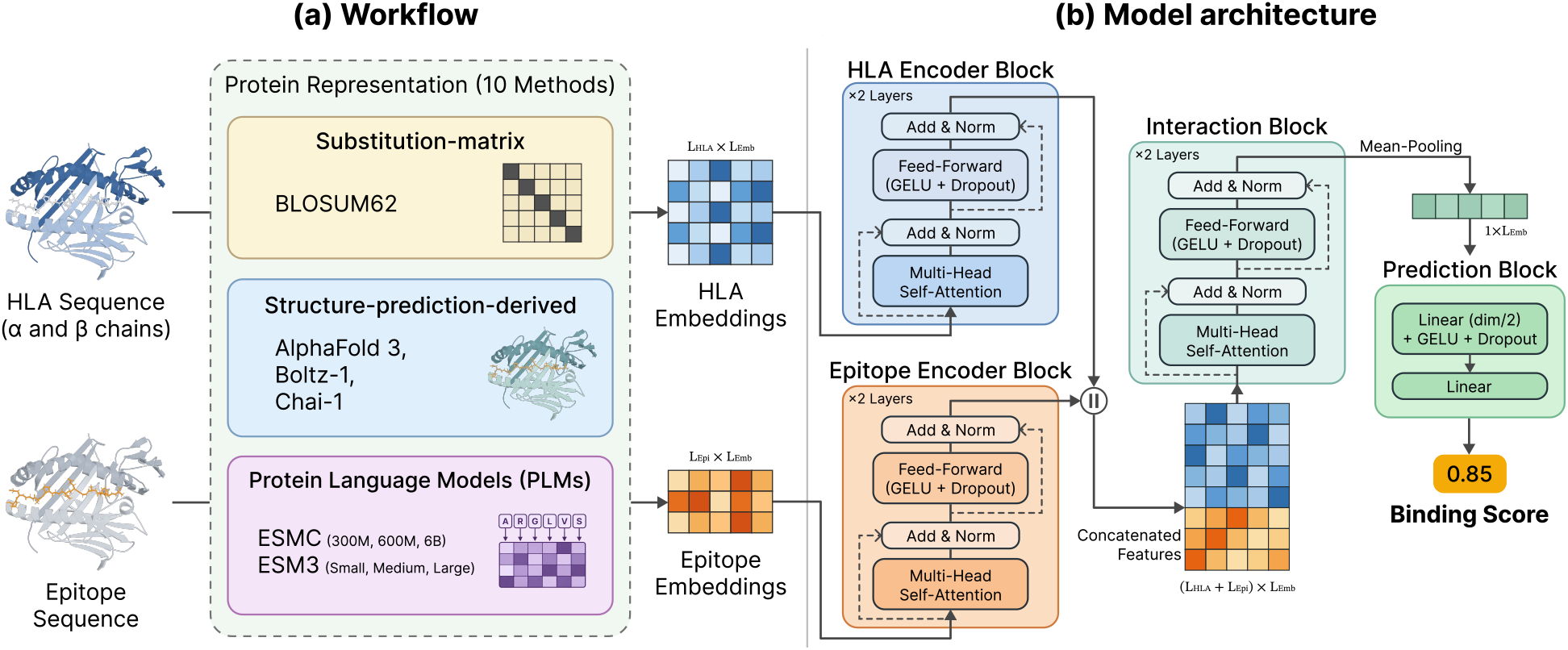
PREpiBind workflow and architecture. (a) Ten representations encode the HLA and epitope inputs. The molecular illustration shows HLA-DR4 bound to a collagen type II peptide (PDB 7NZF) [37]. (b) Separate encoders process HLA and epitope features before concatenation, joint interaction, mean pooling, and binding-score prediction.

#### Training and Evaluation

Model training was performed with a fixed batch size of 128 using the AdamW optimizer. To identify optimal convergence, we evaluated three distinct learning rates (10^*−*3^, 10^*−*4^, and 10^*−*5^) for each model. Based on validation performance, a learning rate of 10^*−*3^ was optimal for the BLOSUM62 and DeepNeo baselines, whereas all other representations achieved peak performance at 10^*−*5^. To prevent overfitting, early stopping was implemented with a patience of 10 epochs, monitoring the validation loss. All experiments used three random initialization seeds (42, 100, and 128), with five folds per seed. No seed was selected. We first averaged the five folds within each seed and then averaged the three seed-level values. Benchmark results are reported as mean *±* standard deviation across the three seed-level means (ddof = 1). We did not treat the 15 fold– seed values as independent because folds share a common test set. The same aggregation was used for allele-wise, LOMO, and H2-out analyses, yielding one value per allele or withheld molecule.

#### Protein Representation

The ten embedding strategies span three paradigms of protein encoding: a classical substitution matrix (BLOSUM62), structure-prediction-derived representations (AlphaFold 3, Boltz-1, and Chai-1), and protein language models at several scales (ESMC and ESM3). For the structure-prediction-derived embeddings, we extracted single-residue and pairwise interaction features from the three folding models and applied dimensionality reduction to create unified representations. Note that the structure-prediction-derived representations were alignment-conditioned on the MHC side: the MHC chains supplied to the structure prediction models carried multiple sequence alignments built against large sequence databases, whereas the BLOSUM62, ESMC and ESM3 representations were derived from the MHC and peptide sequences alone (Supplementary Text). Complete technical specifications, including embedding dimensions and preprocessing procedures for all ten representations, are detailed in Supplementary Text 1.2 and Supplementary Table S5.

### Benchmarking Framework and Evaluation Strategy

#### Performance Metrics

ROC-AUC was used as the primary metric because it is threshold-independent and applicable to both PREpiBind outputs and percentile-rank baselines. PR-AUC, F1, accuracy, and MCC were additionally reported in the Supplementary Tables but were not used to rank representations; their definitions are given in Supplementary Text 1.4. Model logits were transformed to probabilities using the sigmoid function, while DeepNeo probabilities were used directly. F1, accuracy, and MCC were calculated at a probability threshold of 0.5.

Reported standard deviations were calculated across three random seeds and reflect run-to-run training variability. Separately, for the pooled Qualitative benchmark, 95% confidence intervals were estimated from 1,000 bootstrap resamples of the test set, recomputing the folds-then-seeds aggregation for each replicate. Pairwise differences between configurations were evaluated using paired bootstrap resampling (Supplementary Table S11).

ROC-AUC was aggregated two ways: *pooled*, ranking all test peptide–allele pairs on a single score axis (Table 2), and *per-allele*, computed within each allele and averaged across alleles (Figure 3a). Allele-wise ROC-AUC was reported only for alleles with at least five positive and five negative test examples. NetMHCIIpan-4.3 and MixMHC2pred-2.0 yield per-allele-calibrated %Rank values, so their pooled and per-allele results are interpreted separately.

**Table 2.**
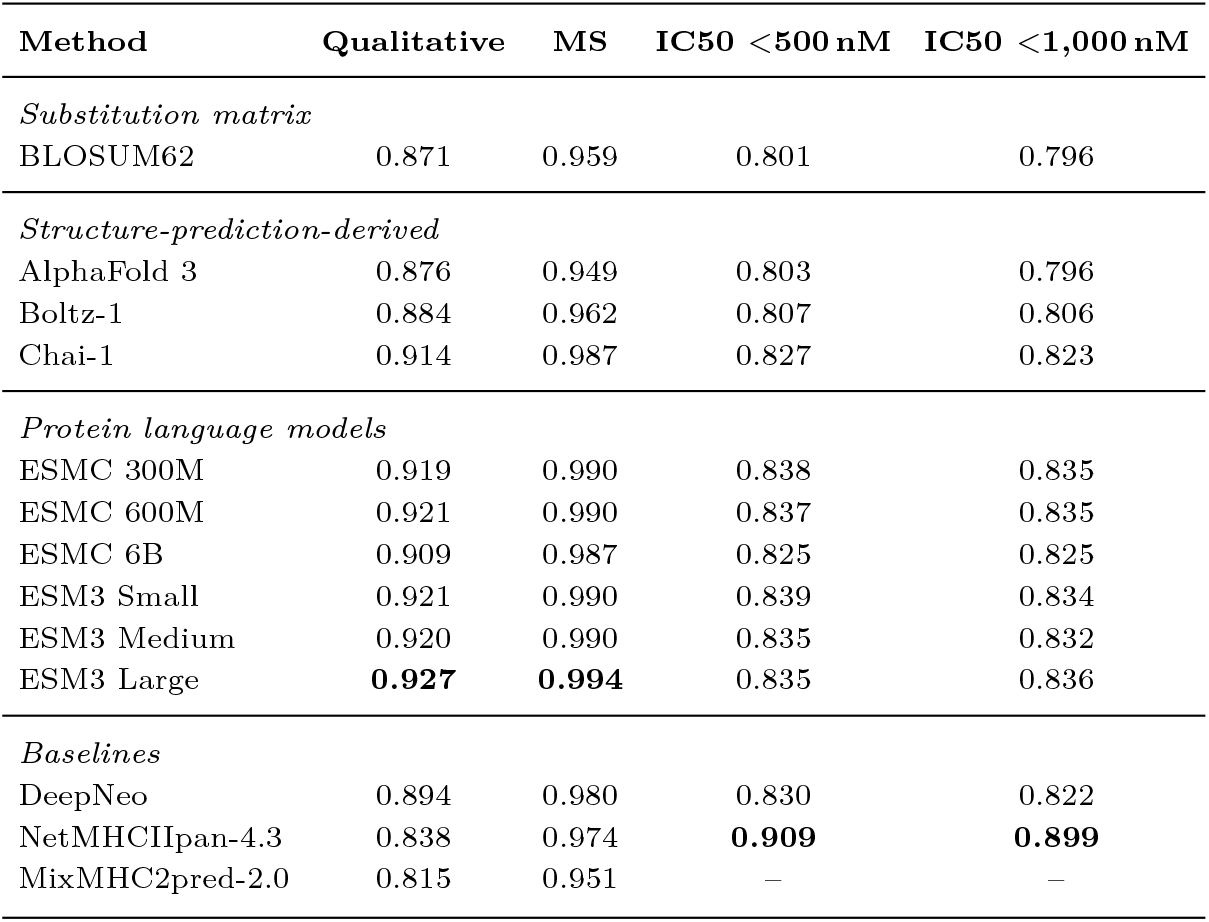
ROC-AUC across all evaluated methods and datasets. Methods are grouped by representation type, and bold marks the best value per column. ‘–’ denotes a setting not compared in the main benchmark. Standard deviations and reference-tool scoring conventions are given in Supplementary Tables S7–S10 and Supplementary Text 1.8.

#### Baseline Models

We benchmarked the performance of PREpiBind against those of three widely used methods for MHC-II binding prediction: DeepNeo [2, 15, 38], NetMHCIIpan-4.3 [14], and MixMHC2pred-2.0 [12].

DeepNeo-v2 is only available as a webserver, so we re-implemented it in PyTorch following the published architecture and training procedure (Supplementary Text 1.8). Because it models only the MHC-II *β* chain, *α*-chain sequences were removed from its datasets and the resulting duplicates eliminated, with epitope–*β*-chain pairs assigned uniquely to training or test. The re-implementation reaches a ROC-AUC of 0.894 *±* 0.002 on our Qualitative test set, close to the 0.882 that the original publication reports for its own MHC-II test set [15].

For NetMHCIIpan-4.3 and MixMHC2pred-2.0, precompiled binaries were executed using default configurations, with the single exception that MixMHC2pred-2.0 was run with the `no_context` option enabled to match our standardized input formats. Neither tool covers every allele in our benchmark, so each was scored on the test rows carrying an allele it supports; excluded alleles and sample sizes are given in Supplementary Text 1.9. Both tools were scored on *−*%Rank, the only quantity comparable across them, and a 0.5 cut has no meaning on a percentile rank, so only threshold-independent metrics (ROC-AUC and PR-AUC) were used for these two tools.

#### Generalization Ability Evaluation

Generalization was assessed first from the distribution of allele-wise ROC-AUC values. Subsequently, we conducted leave-one-molecule-out (LOMO) validation to evaluate performance when the target MHC molecule was withheld during training. LOMO withholds an entire MHC molecule together with all of its peptides. Note that this is a group-wise held-out-molecule analysis and not observation-level leave-one-out cross-validation. LOMO was used neither for checkpoint selection nor as the headline performance estimate, which is reported on the independent test set. Because LOMO requires retraining the model with all data for the target allele withheld, the analysis was restricted to retrainable implementations (all PREpiBind variants and our DeepNeo re-implementation). LOMO was performed on the Qualitative dataset for 47 MHC molecules with at least 20 samples in each class and a minority-class fraction of at least 20% (38 molecules met these criteria for the *β*-chain-only DeepNeo). Within each LOMO iteration, the model was trained using stratified five-fold cross-validation on the remaining data, and performance for the withheld molecule was aggregated over folds and seeds as above.

To evaluate cross-species generalization, we performed a “species-out” experiment by excluding all mouse (H2) data during training and evaluating the models exclusively on H2 alleles. The Wilcoxon signed-rank test was used to compare performance distributions between models, with the allele or withheld molecule as the statistical unit. DeepNeo models no MHC-II *α* chain, so its units are bare *β* chains and share no labels with the *α*/*β* pairs used for the other models. For comparisons involving DeepNeo, the other models’ per-unit ROC-AUC values were therefore averaged within each *β* chain. *p*-values were adjusted across all model pairs within a panel using the Holm–Bonferroni procedure.

## Results

### Benchmark performance across datasets

Across the evaluated datasets, PLM-based representations generally yielded higher predictive scores than the structure-prediction-derived and substitution-matrix representations (Figure 2). On the Qualitative dataset, ESM3 Large yielded the highest observed ROC-AUC (0.927 *±* 0.002). The results with ESMC 600M, ESM3 Small, ESM3 Medium, and ESMC 300M clustered at 0.919–0.921, within their seed-level variation. All evaluated PLM variants exceeded our PyTorch re-implementation of DeepNeo (0.894*±*0.002). The results of structure-prediction-derived representations were heterogeneous: Chai-1 reached 0.914 *±* 0.001, whereas AlphaFold 3 (0.876 *±* 0.006) and Boltz-1 (0.884 *±* 0.004) were close to BLOSUM62 (0.871 *±* 0.005). NetMHCIIpan-4.3 (0.838) and MixMHC2pred-2.0 (0.815) were lower than all PREpiBind representations on this dataset. Pairwise bootstrap comparisons for representative configurations are reported in Supplementary Table S11.

**Fig. 2.**
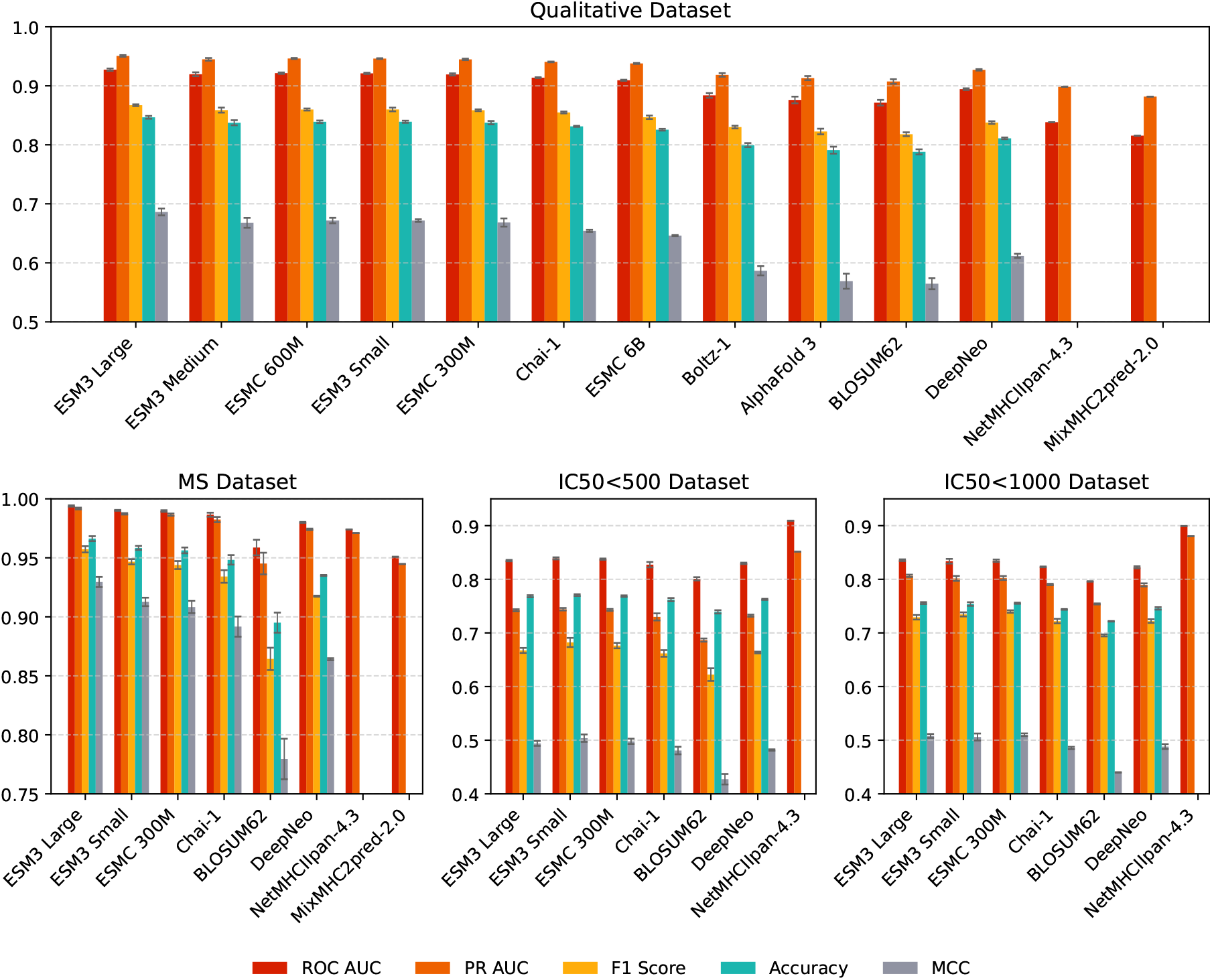
Benchmark performance across Qualitative, MS, IC50*<*500, and IC50*<*1000 datasets. Grouped bars show ROC-AUC, PR-AUC, F1, accuracy, and MCC. Only ROC-AUC and PR-AUC are reported for percentile-rank tools. Complete values are given in Table 2 and Supplementary Tables S7–S10.

Since homologous binding cores are a recognized confounder in pMHC benchmarking [39], we also examined test epitopes sharing no 9-mer with training. This stratum contained 18.6% of unique test epitopes. The ordering of the trained representations was unchanged, while BLOSUM62 lost 0.115 ROC-AUC compared with 0.038–0.045 for Chai-1, ESMC 300M, and ESM3 Small (Supplementary Text 1.5 and Supplementary Tables S2 and S3).

To determine whether differences in the intrinsic separation of MHC-II gene families across representations could explain these performance differences, we projected the MHC-II embeddings with UMAP and quantified their gene-family separation with silhouette coefficients. BLOSUM62 separated the gene families as effectively as Chai-1 (macro-averaged silhouette score, 0.55 versus 0.56) despite showing substantially lower predictive performance (Supplementary Text 1.3, Supplementary Table S12, and Supplementary Figure S2). Thus, separation of MHC-II gene families alone does not explain the observed differences in predictive performance.

On the MS dataset, ESM3 Large again yielded the highest ROC-AUC (0.994 *±* 0.001). Four other PLM configurations scored 0.990, followed by Chai-1 and ESMC 6B at 0.987. Complete baseline values are reported in Table 2.

On the thresholded IC50 datasets, the leading PLM configurations clustered at 0.835–0.839 for 500 nM and 0.832–0.836 for 1,000 nM, with no stable internal ordering. Those of Chai-1 and DeepNeo representations were slightly lower. NetMHCIIpan-4.3 scored 0.909 and 0.899 using its binding-affinity head, which is fitted to measured IC50 values [14]; these values are therefore interpreted separately from the frozen-representation comparisons. MixMHC2pred-2.0 has no affinity head and is not compared in the main IC50 panels; its measured values are reported in Supplementary Tables S9 and S10.

### PREpiBind performance across allele-wise and withheld-molecule evaluations

We next evaluated PREpiBind under allele-wise, LOMO, H2-specific LOMO, and H2-out settings (Figure 3). In the allele-wise analysis, PLM representations also led to the best prediction performance. Chai-1 and both PLM representations outperformed BLOSUM62, whereas ESMC 300M and ESM3 Small showed equivalent performance (Supplementary Table S18). Mean ROC-AUC values over the 57 alleles shared by the pair-named methods were 0.780 for BLOSUM62, 0.843 for Chai-1, and 0.852 for both ESMC 300M and ESM3 Small. DeepNeo averaged 0.811 over its own 48 *β* chains. Complete comparison results including DeepNeo on the shared *β*-chain unit are given in Supplementary Table S19.

**Fig. 3.**
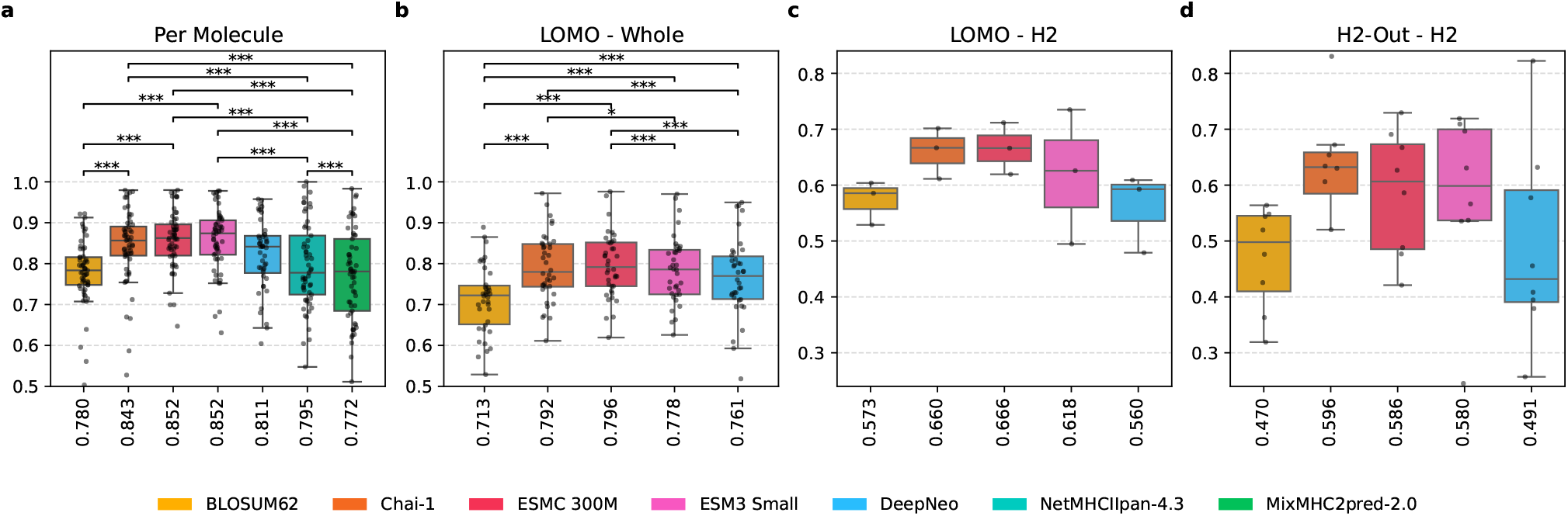
Prediction performance across various evaluation settings. (a) Allele-wise ROC-AUC, (b) LOMO, (c) H2-specific LOMO, and (d) H2-out evaluation. Each point represents one molecule and the number beside each box is the mean. Brackets show Holm-corrected paired comparisons; complete values and omitted comparisons are reported in Supplementary Tables S14–S20. Panels (a,b) and (c,d) use separate y-axis ranges.

NetMHCIIpan-4.3 (0.795) and MixMHC2pred-2.0 (0.772) showed lower per-allele performance than Chai-1 and both PLMs on their supported alleles. Note that these published tools are included for contextual comparison rather than as like-for-like benchmarks (Supplementary Text 1.8).

LOMO directly tested transfer to molecules absent from training (Figure 3b). In the LOMO scenario, ESMC 300M and Chai-1 representations led to comparable prediction performance followed by ESM3 Small, DeepNeo and BLOSUM62. Per-molecule mean ROC-AUC, over the 38 *β*-chain units shared with DeepNeo, was 0.796 for ESMC 300M, 0.792 for Chai-1, 0.778 for ESM3 Small, 0.761 for DeepNeo, and 0.713 for BLOSUM62 (Supplementary Tables S16 and S20). For contextual comparison, NetMHCIIpan-4.3 and MixMHC2pred-2.0 had per-molecule means of 0.805 and 0.779, respectively. However, neither model was retrained under LOMO, and therefore their training data were not controlled to exclude the held-out molecules.

H2-specific LOMO contained only the three H2 molecules meeting the inclusion criteria and was therefore interpreted descriptively (Figure 3c). Chai-1 and the two PLM configurations had higher observed mean AUC values than BLOSUM62 and DeepNeo, but the panel is too small to support a robust ordering.

In H2-out evaluation, models trained only on human data were evaluated on all murine H2 data (3,120 rows across nine molecules). Pooled ROC-AUC was 0.682 for ESM3 Small, 0.680 for ESMC 300M, 0.628 for Chai-1, 0.529 for DeepNeo, and 0.476 for BLOSUM62. When each of the eight molecules with at least five positives and five negatives was weighted equally (Figure 3d), the ordering changed: Chai-1 averaged 0.596, ESMC 300M 0.586, and ESM3 Small 0.580. With only eight molecules, the paired analysis lacked the resolution to establish a stable ordering among these leading representations; the aggregation sensitivity and statistical limitations are reported in Supplementary Text 1.6 and Supplementary Table S16. NetMHCIIpan-4.3 and MixMHC2pred-2.0 scored higher in this setting, but both ship murine H-2 models [12, 14]. H2-out is therefore a cross-species extrapolation only for the retrained representations.

### Serotype-specific performance analysis

We also performed serotype-specific analyses to assess whether the overall performance trends were consistent across MHC-II serotypes. PLM representations generally maintained their performance advantage across serotypes, although absolute performance varied substantially among serotypes, with HLA-DP yielding the highest values in both datasets (Figure 4). On the Qualitative dataset, ESM3 Small, ESMC 300M, and Chai-1 reached 0.983, 0.982, and 0.980 on HLA-DP and remained above BLOSUM62 and DeepNeo across the three human serotypes, although their internal ordering changed by serotype. Murine H2 resulted in the lowest range (0.813– 0.871). Both published tools were higher on H2, but they carry murine models and are therefore interpreted separately. Detailed serotype values and comparisons are given in Supplementary Text 1.6.

**Fig. 4.**
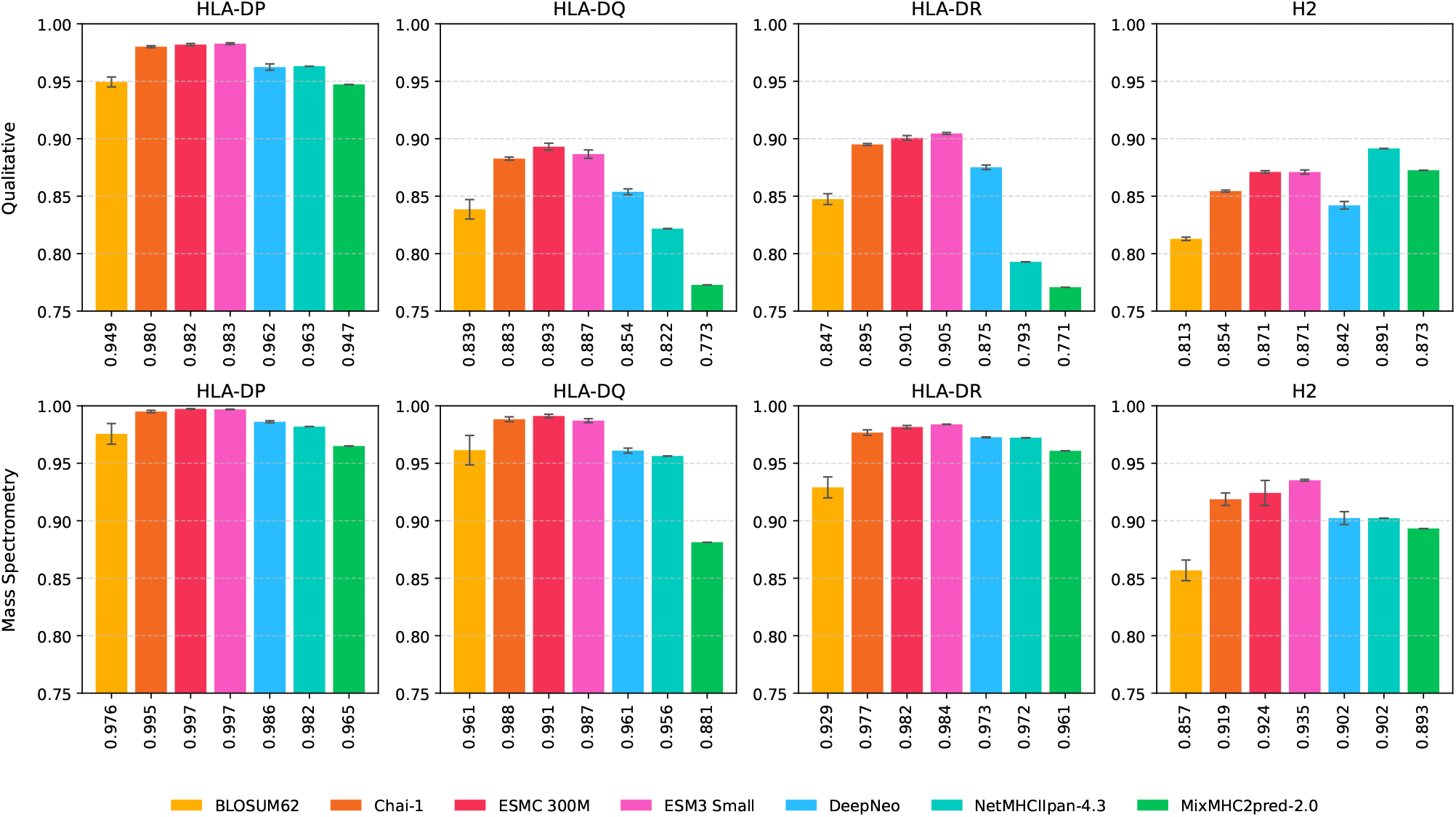
Serotype-level ROC-AUC for HLA-DP, HLA-DQ, HLA-DR, and murine H2. Rows show Qualitative and MS results; columns show serotypes. Labels give ROC-AUC values and whiskers show standard deviations for trained models.

### The relationship between representation model size and prediction accuracy

It is well established that pretraining loss can improve with model scale [40], including protein language models [21, 22]. We tested whether a similar scaling relationship holds for pMHC-II prediction. The results show that overall downstream pMHC-II accuracy appears to increase with representation size (Figure 5), but not consistently. ESM3 Large led to the highest observed value (0.927 *±* 0.002), whereas ESM3 Small and Medium results were slightly lower and indistinguishable at 0.921 *±* 0.001 and 0.920 *±* 0.003. Within ESMC, 300M and 600M model results differed only within seed-level variation (0.919 versus 0.921), while the largest model ESMC 6B resulted in lower performance at 0.909. Among the structure-prediction-derived representations, Chai-1 achieved the highest performance.

**Fig. 5.**
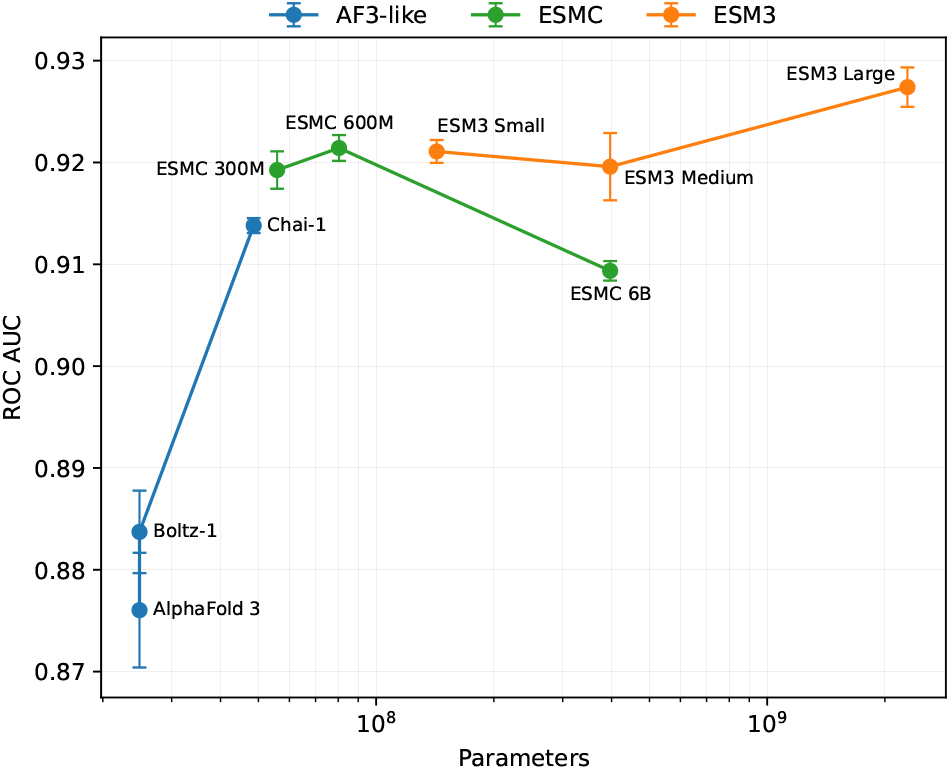
ROC-AUC on the Qualitative test set versus the number of trained PREpiBind parameters (log scale), which is set by the width of the input representation and, within each PLM family, increases with the size of the pretrained model. BLOSUM62 is excluded as it is not a pretrained model. Error bars show the standard deviation across three seeds.

## Discussion

Overall, our benchmark results show that PLM representations yielded the highest pooled performance among the tested representations on the Qualitative, MS, and thresholded IC50 datasets. The benchmark results also demonstrate that their advantage narrows under the molecule-focused and generalization-focused evaluations. Pooled ROC-AUC was also higher than per-allele ROC-AUC for every representation, which is consistent with greater weighting of data-rich alleles (Supplementary Text 1.7 and Supplementary Table S13). The Chai-1 embedding was competitive with the leading PLMs under LOMO, while H2-out evaluation showed that their relative ranking was sensitive to whether performance was aggregated across samples or equally across MHC molecules. These results suggest that the relative advantage of PLM representations may be greater for well-represented MHC molecules, whereas the alignment-conditioned Chai-1 representation may be more competitive under data-sparse or molecule-level generalization settings. Similar task dependence has been reported in other protein-representation benchmarks, where no pretrained model is uniformly best and the adaptation strategy also affects downstream performance [41, 42]. Therefore, representation choice should be guided by the intended application rather than by a single global ranking.

Our results further show substantial variation among representations derived from the three structure-prediction models, indicating that these representations should not be treated as a homogeneous class. Importantly, our benchmark evaluates their transferability through the residue-level PREpiBind interface rather than the intrinsic information content of the pretrained models. Pairwise representations were reduced to sequence-aligned features, and MHC and epitope embeddings were generated independently; thus, cross-interface pair information was not directly provided to PREpiBind. Alternative interfaces that retain pairwise or structural information more explicitly may yield different relative performance [43–46]. In addition, whereas PLM embeddings were generated from single sequences, the structure-prediction-derived MHC representations incorporated alignment information through their respective model trunks.

There are several limitations that should be addressed in future studies. The framework is restricted to 15-mer peptides, which capture the canonical 9-mer core with its flanking residues but leave length-agnostic training to future work. The assay-derived settings overlap substantially, and the MS benchmark additionally draws its non-binders from the Qualitative dataset. Its high values may therefore reflect systematic differences between assay sources as well as pMHC signal and should not be read as an independent replication. Every representation was used as a frozen feature extractor with one fixed downstream architecture. Thus, the results describe transfer under that protocol rather than the best attainable performance for each representation [42]. Related PLM-based pMHC-II predictors were not included in the head-to-head comparison because they use different adaptation strategies and benchmark settings. For example, pMHChat fine-tunes its encoders and uses a different dataset and peptide-length range [27]. A direct comparison would require retraining under the same splits. Whether PLMs encode information about alternative or reverse binding modes, and whether they complement the specialized features of NetMHCIIpan-4.3 and MixMHC2pred-2.0, remains to be tested with dedicated motif and binding-core analyses. Finally, clinical immunogenicity depends on intracellular processing efficiency and T-cell receptor recognition beyond the binding step modelled here [47, 48].

### Conclusion

In this study, we present PREpiBind, a joint-attention framework for systematically evaluating diverse protein representations for pMHC-II binding prediction. Under a common downstream architecture and identical data splits, PLM representations, particularly ESM3 variants, achieved the strongest overall performance in pooled evaluations. However, their advantage narrowed under molecule-focused generalization settings, with Chai-1 becoming competitive under LOMO and H2-out evaluation showing sensitivity to the aggregation scheme. These findings highlight that representation performance depends on the intended prediction setting and should not be summarized by a single global ranking. Broader allele panels, additional species, and independently curated external benchmarks will be important for establishing the generality of these observations. By releasing PREpiBind as a modular, open-source framework, we aim to facilitate reproducible evaluation and integration of emerging protein representations for pMHC-II prediction and related computational immunology applications.

## Data Availability

Source code, the training and evaluation datasets, and the machine-readable result tables D01–D12 are available at https://github.com/daylight-00/PREpiBind [49]; the code is released under the MIT License. The version submitted here is archived at Zenodo under https://doi.org/10.5281/zenodo.22934281. Trained model checkpoints are deposited on HuggingFace at https://huggingface.co/daylight-00/prepibind, with half-precision copies for the demonstration notebooks at https://huggingface.co/daylight-00/prepibind-demo, and pre-computed MHC class II embeddings at https://huggingface.co/datasets/daylight-00/prepibind-embeddings [50]. The ESMC 300M base model weights originate with Chan Zuckerberg Biohub (https://huggingface.co/biohub/esmc-300m-2024-12) and are released under the MIT License; the copy used here is mirrored at https://huggingface.co/daylight-00/esmc-300m-2024-12. Every quantitative figure and table in this work can be regenerated from the repository alone. The raw model predictions from which every reported number is computed, together with the IEDB export the datasets were built from, are archived at Zenodo under https://doi.org/10.5281/zenodo.22857373 [51]. Peptide–MHC binding data were obtained from the Immune Epitope Database (IEDB) [29] (https://www.iedb.org, Database Export v3, build of 2025/04/21), which distributes its data under a Creative Commons Attribution 4.0 International licence; the datasets redistributed here are modified from that export, having been filtered, relabelled and re-split, and attribution to the submitting authors is carried by the named snapshot. MHC allele sequences were retrieved from the IPD-IMGT/HLA Database [30–32], release 3.59.0 (2025-01-15), obtained from its public distribution at https://github.com/ANHIG/IMGTHLA, and murine H2 sequences from UniProt [33] (https://www.uniprot.org/). The IPD-IMGT/HLA-derived sequences redistributed with this work are published by permission of Anthony Nolan.

## Author contributions statement

D.H.J.: Conceptualization, Investigation, Methodology, Data curation, Formal analysis, Software, Validation, Visualization, Funding acquisition, Writing – Original draft, Writing – Review & Editing. D.K.: Validation, Project administration, Visualization, Writing – Review & Editing. U.H.: Investigation, Data curation, Funding acquisition. B.P.: Investigation, Data curation, Funding acquisition. Y.C.: Validation, Supervision, Funding acquisition, Writing – Review & Editing. J.L.: Conceptualization, Supervision, Resources, Funding acquisition, Writing – Review & Editing.

## Acknowledgments

The authors thank all members of the Laboratory of Computational Drug Discovery at Seoul National University for their helpful discussions and feedback throughout this work.

Generative AI tools were used for language editing and refinement of author-prepared drafts, assistance with validation of analysis code, and consistency checks among the manuscript, supplementary materials, and underlying data. All outputs were reviewed and verified by the authors, who take full responsibility for the final content, analyses, and conclusions (Supplementary Text 1.10).

## References

1. Paul A Roche and Kazuyuki Furuta. The ins and outs of MHC class II-mediated antigen processing and presentation. Nature Reviews Immunology, 15(4):203–216, 2015.

2. Jeong Yeon Kim, Hongui Cha, Kyeonghui Kim, Changhwan Sung, Jinhyeon An, Hyoeun Bang, Hyungjoo Kim, Jin Ok Yang, Suhwan Chang, Incheol Shin, et al. MHC II immunogenicity shapes the neoepitope landscape in human tumors. Nature Genetics, 55(2):221–231, 2023.

3. Irina A Ishina, Maria Y Zakharova, Inna N Kurbatskaia, Azad E Mamedov, Alexey A Belogurov Jr, and Alexander G Gabibov. MHC class II presentation in autoimmunity. Cells, 12(2):314, 2023.

4. Elise Alspach, Danielle M Lussier, Alexander P Miceli, Ilya Kizhvatov, Michel DuPage, Adrienne M Luoma, Wei Meng, Cheryl F Lichti, Ekaterina Esaulova, Anthony N Vomund, et al. MHC-II neoantigens shape tumour immunity and response to immunotherapy. Nature, 574(7780):696–701, 2019.

5. Massimo Andreatta, Edita Karosiene, Michael Rasmussen, Anette Stryhn, Søren Buus, and Morten Nielsen. Accurate pan-specific prediction of peptide-MHC class II binding affinity with improved binding core identification. Immunogenetics, 67(11–12):641–650, 2015.

6. Massimo Andreatta, Vanessa I Jurtz, Thomas Kaever, Alessandro Sette, Bjoern Peters, and Morten Nielsen. Machine learning reveals a non-canonical mode of peptide binding to MHC class II molecules. Immunology, 152(3):255–264, 2017.

7. Christopher J Holland, David K Cole, and Andrew Godkin. Redirecting CD4+ T cell responses with the flanking residues of MHC class II-bound peptides: the core is not enough. Frontiers in Immunology, 4:172, 2013.

8. Bruce J MacLachlan, Garry Dolton, Athanasios Papakyriakou, Alexander Greenshields-Watson, Georgina H Mason, Andrea Schauenburg, Matthieu Besneux, Barbara Szomolay, Tim Elliott, Andrew K Sewell, et al. HLA class II peptide flanking residues tune the immunogenicity of a human tumor-derived epitope. Journal of Biological Chemistry, 294(52):20246–20258, 2019.

9. Calliope A Dendrou, Jan Petersen, Jamie Rossjohn, and Lars Fugger. HLA variation and disease. Nature Reviews Immunology, 18(5):325–339, 2018.

10. Dominic J Barker, Giuseppe Maccari, Xenia Georgiou, Michael A Cooper, Paul Flicek, James Robinson, and Steven GE Marsh. The IPD-IMGT/HLA database. Nucleic Acids Research, 51(D1):D1053–D1060, 2023.

11. Daniel T Rademaker, Farzaneh M Parizi, Marieke van Vreeswijk, Sanna Eerden, Dario F Marzella, and Li C Xue. Predicting reverse-bound peptide conformations in MHC class II with PANDORA. Frontiers in Immunology, 16:1525576, 2025.

12. Julien Racle, Philippe Guillaume, Julien Schmidt, Justine Michaux, Amédé Larabi Kelvin Lau, Marta AS Perez, Giancarlo Croce, Raphaël Genolet, George Coukos, et al. Machine learning predictions of MHC-II specificities reveal alternative binding mode of class II epitopes. Immunity, 56(6):1359–1375, 2023.

13. Steven Henikoff and Jorja G Henikoff. Amino acid substitution matrices from protein blocks. Proceedings of the National Academy of Sciences, 89(22):10915–10919, 1992.

14. Jonas B Nilsson, Saghar Kaabinejadian, Hooman Yari, Michel GD Kester, Peter van Balen, William H Hildebrand, and Morten Nielsen. Accurate prediction of HLA class II antigen presentation across all loci using tailored data acquisition and refined machine learning. Science Advances, 9(47):eadj6367, 2023.

15. Jeong Yeon Kim, Hyoeun Bang, Seung-Jae Noh, and Jung Kyoon Choi. DeepNeo: a webserver for predicting immunogenic neoantigens. Nucleic Acids Research, 51(W1):W134–W140, 2023.

16. Peiyuan Feng, Jianyang Zeng, and Jianzhu Ma. Predicting MHC-peptide binding affinity by differential boundary tree. Bioinformatics, 37(Supplement_1):i254–i261, 2021.

17. Ronghui You, Wei Qu, Hiroshi Mamitsuka, and Shanfeng Zhu. DeepMHCII: a novel binding core-aware deep interaction model for accurate MHC-II peptide binding affinity prediction. Bioinformatics, 38(Supplement_1):i220–i228, 2022.

18. Eric Glynn, Dario Ghersi, and Mona Singh. Towards equitable MHC binding predictions: computational strategies to assess and reduce data bias. bioRxiv, 2024.

19. Jia-Ying Chen, Jing-Fu Wang, Yue Hu, Xin-Hui Li, Yu-Rong Qian, and Chao-Lin Song. Evaluating the advancements in protein language models for encoding strategies in protein function prediction: a comprehensive review. Frontiers in Bioengineering and Biotechnology, 13:1506508, 2025.

20. Yajie Meng, Zhuang Zhang, Chang Zhou, Xianfang Tang, Xinrong Hu, Geng Tian, Jialiang Yang, and Yuhua Yao. Protein structure prediction via deep learning: an in-depth review. Frontiers in Pharmacology, 16:1498662, 2025.

21. Thomas Hayes, Roshan Rao, Halil Akin, Nicholas J Sofroniew, Deniz Oktay, Zeming Lin, Robert Verkuil, Vincent Q Tran, Jonathan Deaton, Marius Wiggert, et al. Simulating 500 million years of evolution with a language model. Science, page eads0018, 2025.

22. Salvatore Candido, Thomas Hayes, Alexander Derry, Roshan Rao, Zeming Lin, Robert Verkuil, Bryan Z Wu, Jin Sub Lee, Elise S Bruguera, Jehan A Keval, et al. Language modeling materializes a world model of protein biology. bioRxiv, page 2026.06.03.729735, 2026.

23. Josh Abramson, Jonas Adler, Jack Dunger, Richard Evans, Tim Green, Alexander Pritzel, Olaf Ronneberger, Lindsay Willmore, Andrew J Ballard, Joshua Bambrick, et al. Accurate structure prediction of biomolecular interactions with AlphaFold 3. Nature, 630(8016):493–500, 2024.

24. Chai Discovery Team, Jacques Boitreaud, Jack Dent, Matthew McPartlon, Joshua Meier, Vinicius Reis, Alex Rogozhonikov, and Kevin Wu. Chai-1: Decoding the molecular interactions of life. bioRxiv, 2024.

25. Jeremy Wohlwend, Gabriele Corso, Saro Passaro, Mateo Reveiz, Ken Leidal, Wojtek Swiderski, Tally Portnoi, Itamar Chinn, Jacob Silterra, Tommi Jaakkola, et al. Boltz-1: Democratizing biomolecular interaction modeling. bioRxiv, 2024.

26. Zeming Lin, Halil Akin, Roshan Rao, Brian Hie, Zhongkai Zhu, Wenting Lu, Nikita Smetanin, Robert Verkuil, Ori Kabeli, Yaniv Shmueli, Allan dos Santos Costa, Maryam Fazel-Zarandi, Tom Sercu, Salvatore Candido, and Alexander Rives. Evolutionary-scale prediction of atomic-level protein structure with a language model. Science, 379(6637):1123–1130, 2023.

27. Jiani Ma, Zhikang Wang, Cen Tong, Qi Yang, Lin Zhang, and Hui Liu. pMHChat, characterizing the interactions between major histocompatibility complex class II molecules and peptides with large language models and deep hypergraph learning. Briefings in Bioinformatics, 26(4):bbaf321, 2025.

28. Soobon Ko, Honglan Li, Hongeun Kim, Woong-Hee Shin, Junsu Ko, and Yoonjoo Choi. Benchmarking sequence-based and AlphaFold-based methods for pMHC-II binding core prediction: Distinct strengths and consensus approaches. bioRxiv, 2024.

29. Randi Vita, Nina Blazeska, Daniel Marrama, IEDB Curation Team, Sebastian Duesing, Jason Bennett, Jason Greenbaum, Marcus De Almeida Mendes, Jarjapu Mahita, Daniel K Wheeler, et al. The immune epitope database (IEDB): 2024 update. Nucleic Acids Research, 53(D1):D436–D443, 2025.

30. Dominic J Barker, Richard H L Natarajan, Michael A Cooper, Sebastian J F Hopper, Andrew D Yates, Peter Parham, Steven G E Marsh, and James Robinson. The IPD-IMGT/HLA database: recent developments in sequence submission. Nucleic Acids Research, 54(D1):D1152–D1158, 2026.

31. James Robinson, Dominic J Barker, and Steven GE Marsh. 25 years of the IPD-IMGT/HLA database. HLA, 103(6):e15549, 2024.

32. James Robinson, A Malik, Peter Parham, Julia G Bodmer, and Steven GE Marsh. IMGT/HLA database–a sequence database for the human major histocompatibility complex. Tissue antigens, 55(3):280–287, 2000.

33. The UniProt Consortium. UniProt: the universal protein knowledgebase in 2025. Nucleic Acids Research, 53(D1):D609–D617, 2025.

34. Typhaine Paysan-Lafosse, Matthias Blum, Sara Chuguransky, Tiago Grego, Beatriz Lazaro Pinto, Gustavo A Salazar, Maxwell L Bileschi, Peer Bork, Alan Bridge, Lucy Colwell, et al. InterPro in 2022. Nucleic Acids Research, 51(D1):D418–D427, 2023.

35. Weilong Zhao and Xinwei Sher. Systematically benchmarking peptide-MHC binding predictors: From synthetic to naturally processed epitopes. PLoS Computational Biology, 14(11):e1006457, 2018.

36. Scott Southwood, John Sidney, Akihiro Kondo, Marie-France del Guercio, Ettore Appella, Stephen Hoffman, Ralph T Kubo, Robert W Chesnut, Howard M Grey, and Alessandro Sette. Several common HLA-DR types share largely overlapping peptide binding repertoires. The Journal of Immunology, 160(7):3363–3373, 1998.

37. Changrong Ge, Sebastian Weisse, Bingze Xu, Doreen Dobritzsch, Johan Viljanen, Jan Kihlberg, Nhu-Nguyen Do, Nils Schneider, Harald Lanig, Rikard Holmdahl, and Harald Burkhardt. Key interactions in the trimolecular complex consisting of the rheumatoid arthritis-associated DRB1*04:01 molecule, the major glycosylated collagen II peptide and the T-cell receptor. Annals of the Rheumatic Diseases, 81(4):480–489, 2022.

38. Kwoneel Kim, Hong Sook Kim, Jeong Yeon Kim, Hyunchul Jung, Jong-Mu Sun, Jin Seok Ahn, Myung-Ju Ahn, Keunchil Park, Se-Hoon Lee, and Jung Kyoon Choi. Predicting clinical benefit of immunotherapy by antigenic or functional mutations affecting tumour immunogenicity. Nature Communications, 11(1):951, 2020.

39. Yohan Kim, John Sidney, Clemencia Pinilla, Alessandro Sette, and Bjoern Peters. Dataset size and composition impact the reliability of performance benchmarks for peptide–MHC binding predictions. BMC Bioinformatics, 15:241, 2014.

40. Jared Kaplan, Sam McCandlish, Tom Henighan, Tom B. Brown, Benjamin Chess, Rewon Child, Scott Gray, Alec Radford, Jeffrey Wu, and Dario Amodei. Scaling laws for neural language models, 2020.

41. Young Su Ko, Jonathan Parkinson, and Wei Wang. Scalable embedding fusion with protein language models: insights from benchmarking text-integrated representations. Briefings in Bioinformatics, 27(1):bbag014, 2026.

42. Thomas Bikias, Evangelos Stamkopoulos, and Sai T Reddy. PLMFit: benchmarking transfer learning with protein language models for protein engineering. Briefings in Bioinformatics, 26(4):bbaf381, 2025.

43. Artem Gazizov, Anna Lian, Casper Goverde, Jonathan Mou, Sergey Ovchinnikov, and Nicholas F. Polizzi. AF2BIND: predicting small-molecule binding sites using the pair representation of AlphaFold2. Nature Methods, 23(3):626–635, 2026.

44. Amir Motmaen, Justas Dauparas, Minkyung Baek, Mohamad H. Abedi, David Baker, and Philip Bradley. Peptide-binding specificity prediction using fine-tuned protein structure prediction networks. Proceedings of the National Academy of Sciences, 120(9):e2216697120, 2023.

45. Jin Su, Chenchen Han, Yuyang Zhou, Junjie Shan, Xibin Zhou, and Fajie Yuan. SaProt: Protein language modeling with structure-aware vocabulary. In The Twelfth International Conference on Learning Representations (ICLR), 2024.

46. Yo Akiyama, Zhidian Zhang, Olivia Tang, Rachel Seongeun Kim, Milot Mirdita, Martin Steinegger, and Sergey Ovchinnikov. Expanding the scope of protein language modeling to protein–protein interactions with MSA Pairformer. Cell, 189(16):4964–4979.e8, 2026.

47. Jacques Neefjes, Marlieke LM Jongsma, Petra Paul, and Oddmund Bakke. Towards a systems understanding of MHC class I and MHC class II antigen presentation. Nature Reviews Immunology, 11(12):823–836, 2011.

48. Markus G Rudolph, Robyn L Stanfield, and Ian A Wilson. How TCRs bind MHCs, peptides, and coreceptors. Annu. Rev. Immunol., 24(1):419–466, 2006.

49. David Hyunyoo Jang, Dongwoo Kim, Untaek Hwang, Byungho Park, Yoonjoo Choi, and Juyong Lee. [software] PREpiBind: source code, training and evaluation datasets, and trained model checkpoints. Zenodo, 2026. Version 1.0.1, archived from https://github.com/daylight-00/PREpiBind. 10.5281/zenodo.22934281.

50. David Hyunyoo Jang, Dongwoo Kim, Untaek Hwang, Byungho Park, Yoonjoo Choi, and Juyong Lee. [dataset] PREpiBind pre-computed MHC class II embeddings. Hugging Face, 2026. https://huggingface.co/datasets/daylight-00/prepibind-embeddings.

51. David Hyunyoo Jang, Dongwoo Kim, Untaek Hwang, Byungho Park, Yoonjoo Choi, and Juyong Lee. [dataset] PREpiBind: prediction snapshot and source data for “Protein Representation-integrated Epitope–MHC Class II Binding Prediction”. Zenodo, 2026. 10.5281/zenodo.22857373.

