## Supplementary Material for "PREpiBind: Protein Representation-integrated Epitope–MHC Class II Binding Prediction"

#### 1 Supplementary Text

##### 1.1 Detailed Dataset Processing and Filtering

###### 1.1.1 IEDB Data Extraction and Preprocessing

All pMHC binding data were retrieved from the Immune Epitope Database (IEDB) Export v3 (dated 2025/04/21). We retained only linear peptides without mutations, and included entries restricted to human and mouse MHC class II alleles. Specifically, we filtered for HLA-DP, HLA-DQ, and HLA-DR alleles annotated to the second field (e.g., HLA-DRB1*01:01), and murine alleles (H2-IAb, IAd, IAg7, IAk, IAq, IAs, IAu, IEd, IEk) with full resolution. This resulted in 977,144 total entries (including duplicates), all subject to uniform preprocessing.

For human MHC class II alleles, we used the second-field resolution to define unique proteins. When multiple alleles shared identical protein sequences within the binding domain, we selected the variant with the longest available sequence to represent the group. Two exceptions—HLA-DPA1*03:06 and HLA-DRB1*01:04—had complementary fragments across variants, which were merged to form consensus sequences.

We filtered out an 8-residue proline-rich insert (PQGPPPAG) from 48 HLA-DQ $\beta$-chains, previously reported to be associated with signaling-related motifs. This insert is not located in the peptide-binding groove and may confound downstream modeling.

Further allele-specific filtering was applied. For HLA-DP and DQ, we required both alpha and beta chains to be specified. HLA-DR entries were retained regardless of alpha chain due to the fixed HLA-DRA*01:01. Duplicated entries from overlapping assays or publications were merged. When conflicting labels were present, we assigned a positive label if any assay reported binding, avoiding resolution of conflicting records as negative.

###### 1.1.2 MS Negative Augmentation

Mass spectrometry-based ligand elution assays identify peptides that are naturally processed and presented by MHC molecules and therefore provide positive examples without explicit negative controls. We constructed a binary classification task by incorporating experimentally annotated negatives from the Qualitative dataset. Published MS-based tools also require negative augmentation, but commonly use sampled decoy peptides rather than the assay-derived negatives used here [1, 2]. Data leakage was prevented by the same $\beta$-chain–peptide grouping applied to the other datasets.

##### 1.1.3 Dataset Composition and Train/Test Splitting

Each dataset (Qualitative, IC50, and MS) was divided into training and test sets with a 70%:30% split. Stratified sampling was applied to preserve the distributions of both class labels (positive/negative) and HLA serotypes in each split. For the IC50 dataset, additional stratification was performed according to binding affinity thresholds: $<$500 nM, 500–1000 nM, and $\geq$1000 nM.

To prevent data leakage and overfitting, all entries with the same $\beta$-chain and peptide sequence were grouped and assigned exclusively to either the training or test set. For models that omit the $\alpha$-chain, we further removed redundant entries that differ only by the presence of $\alpha$-chain annotations.

#### 1.2 Detailed Protein Representation Methods

##### 1.2.1 Downstream Architecture

Each encoder layer comprises a multi-head self-attention sublayer followed by a feed-forward network with GELU activation and dropout, with residual connections and layer normalization applied after each sublayer. The interaction block is a single layer of the same structure. In the prediction head the first linear layer halves the feature dimensionality and the second outputs a scalar. Parameter counts for every representation are given in Supplementary Table S6.

##### 1.2.2 Protein Language Model Embeddings

PLM embeddings were generated from two model series: ESMC and ESM3. Although both build upon ESM-2, their design goals differ: ESM3 emphasizes generative capabilities with integrated structural and contact-map supervision, while ESMC focuses on biologically grounded sequence representations.

For local inference, we used ESMC 300M and 600M models with FlashAttention v2.7.4.post1. For larger models, including ESM3 Small/Medium/Large and ESMC 6B, we accessed the Forge API provided by EvolutionaryScale. All embeddings were generated on NVIDIA H100 SXM5 80GB GPUs and stored in HDF5 format.

##### 1.2.3 Structure-Prediction-Derived Embeddings

For structure-prediction-derived embeddings, we employed outputs from three recent models—AlphaFold 3, Boltz-1, and Chai-1. From each, we extracted the single-residue embedding ($[L, d_\text{s}]$) and the pairwise interaction tensor ($[L, L, d_\text{p}]$). The pairwise tensor was reduced along both residue axes by averaging (mean(0) and mean(1)), yielding two $[L, d_\text{p}]$ matrices. These were concatenated to the single-residue embedding to form a final embedding of size $[L, d_\text{s} + 2d_\text{p}]$.

Since the original implementations did not support direct extraction of features during inference, we modified the source code to expose intermediate representations and bypass diffusion modules. Modified versions are publicly available with traceable commit history.

##### 1.2.4 Traditional Baseline

BLOSUM62 embeddings encode amino acids via substitution scores, providing a low-dimensional reference point despite limited contextual capacity.

##### 1.2.5 MHC Sequence Retrieval and Domain Processing

Since IEDB does not provide MHC protein sequences, we retrieved all allele-specific sequences from the IPD-IMGT/HLA database and UniProt. The IPD-IMGT/HLA database follows the standard four-field HLA nomenclature, where the first and second fields correspond to serotype and protein-coding differences, respectively.

Mouse (H2) MHC class II protein sequences were obtained from UniProt, using curated entries for each allele and chain. For H2-IAg7, the $\alpha$- and $\beta$-chain sequences were confirmed with reference to the structure in PDB ID 6BLX. Alleles annotated with suffixes such as 'N', 'L', 'S', or 'Q', which indicate null or questionable expression, were retained only after removing the suffix to focus on canonical amino acid sequences. In total, we curated $\alpha$- and $\beta$-chain sequences for 7,267 distinct MHC class II alleles.

To isolate the peptide-binding regions of MHC class II molecules, we extracted InterPro-annotated domains from UniProt entries corresponding to each gene. These domains represent the canonical antigen-binding groove and provide consistent structural boundaries across alleles. For each allele, domain boundaries were applied to truncate protein sequences obtained from the IPD-IMGT/HLA database (for human) and UniProt (for mouse).

Embeddings were generated from the full-length protein sequences and trimmed to the annotated binding domains during model input assembly. This preserves the sequence context used for representation generation while restricting the downstream predictor to the peptide-binding domains.

#### 1.3 UMAP visualization of MHC embeddings

To investigate potential intrinsic biases in distinguishing MHC gene families across representations, we visualized the organization of MHC embeddings generated by representative methods (Figure S2). We extracted embeddings for all unique MHC $\alpha$- and $\beta$-chain sequences using the BLOSUM62 baseline, the structure-prediction-derived Chai-1 model, and two PLMs (ESMC 300M and ESM3 Small). After mean-pooling the embeddings across sequence length to generate fixed-dimensional vectors, we performed dimensionality reduction using uniform manifold approximation and projection (UMAP) [3]. To quantify cluster separation, we computed silhouette coefficients (scikit-learn) using cosine distance over the per-allele mean-pooled vectors, with MHC-II gene family as the cluster label as follows:

$$s(i) = \frac{b(i) - a(i)}{\max(a(i), b(i))} \tag{1}$$

, where $a(i)$ is the mean intra-cluster distance and $b(i)$ is the mean nearest-cluster distance. The coefficient ranges from -1 to 1, with values near 1 indicating clear separation between clusters. Gene families represented by at least two alleles were retained (eight families, 113 alleles), and we report the unweighted mean of the per-family silhouette values so that the summary is independent of family size.

The visualizations showed differing degrees of separation among several MHC gene families across representation methods. The visualizations of BLOSUM62, Chai-1, ESMC 300M, and ESM3 Small showed some degree of separation among the displayed gene families, with visually different cluster arrangements across methods. Murine H2 alleles were visually separated from human HLA alleles in all displayed projections. In a quantitative analysis, all four representations yielded comparable positive silhouette values (BLOSUM62, 0.55; Chai-1, 0.56; ESMC 300M, 0.51; ESM3 Small, 0.53). BLOSUM62 separated the displayed MHC-II gene families to a similar degree as Chai-1, ESMC 300M, and ESM3 Small, although their binding-prediction performance differed (Table S12). Gene-family separation alone therefore does not account for the observed differences in pMHC-II prediction performance.

##### 1.4 Performance Metrics Definitions

**ROC-AUC** (Receiver Operating Characteristic Area Under Curve) quantifies the model's ability to distinguish between positive and negative classes over all possible thresholds. It is computed as the area under the ROC curve, which plots the true positive rate (TPR) against the false positive rate (FPR).

**PR-AUC** (Precision-Recall Area Under Curve) is more sensitive to class imbalance and reflects the trade-off between precision and recall across varying thresholds. It is particularly informative when the positive class is underrepresented.

**F1 score** is the harmonic mean of precision and recall, offering a balanced measure of classification performance when both false positives and false negatives are of concern.

**Accuracy** represents the proportion of correctly predicted instances (positive and negative) over the total number of predictions.

**Matthews Correlation Coefficient (MCC)** summarizes the confusion matrix into a single value ranging from $-1$ (complete disagreement) to $+1$ (perfect prediction), with 0 indicating random guessing. MCC is robust to class imbalance and is especially informative in binary classification tasks.

The following equations define each metric in terms of true positives (TP), true negatives (TN), false positives (FP), and false negatives (FN):

$$\text{Accuracy} = \frac{TP + TN}{TP + TN + FP + FN}$$

$$\text{F1 Score} = \frac{2 \cdot TP}{2TP + FP + FN}$$

$$\text{MCC} = \frac{TP \cdot TN - FP \cdot FN}{\sqrt{(TP + FP)(TP + FN)(TN + FP)(TN + FN)}}$$

##### 1.5 Peptide Sequence Redundancy and Train–Test Identity Analysis

###### 1.5.1 Background and Scope of Analysis

A recognised concern in pMHC benchmarking is that two peptides sharing the same 9-mer binding core—but differing in flanking residues—may be assigned to different cross-validation folds or train/test partitions, allowing a model to exploit learned binding-core preferences rather than generalising [4]. NetMHCIIpan addresses this via a common-motif partitioning algorithm that places peptides sharing any 9-mer subsequence into the same cross-validation fold [1, 5].

In PREpiBind, the primary deduplication step groups all entries with the same ($\beta$-chain, peptide) pair and assigns the entire group to either the training or test set exclusively, preventing any identical pMHC binding context from spanning both partitions. This exact-pair grouping does not, however, prevent two peptides sharing a 9-mer subsequence from appearing on opposite sides of the split when they are paired with different alleles—a condition that arises naturally in IEDB data, where the same peptide sequence is routinely tested against multiple alleles. The analysis below quantifies the extent of this overlap and its empirical impact on the reported performance metrics.

Under common-motif partitioning, held-out folds are free of shared 9-mers with the corresponding training folds by construction. Our no-9-mer-overlap stratum is therefore the closest comparison to that evaluation condition. Published approaches generally partition or downweight redundant sequences rather than discarding them from training [1, 2, 4].

##### 1.5.2 9-mer Subsequence Overlap Between Training and Test Epitopes

For each unique 15-mer test epitope we extracted all seven possible 9-mer subsequences and determined whether any of them appeared among the 9-mers derived from unique training epitopes. For each test epitope, the maximum 15-mer pairwise identity to any training epitope was also computed. Results for the Qualitative dataset are shown in Table S2.

The high exact-match rate (58.5%) reflects the IEDB experimental design: the same peptide is frequently tested against multiple alleles, so the same 15-mer sequence appears in both training and test sets when paired with different alleles. The ($\beta$-chain, peptide) grouping prevents exact pMHC-pair leakage; however, 23.0% of test epitopes share a 9-mer with training without being exact matches—the category most relevant to binding-core leakage.

##### 1.5.3 Stratified Performance Analysis

We stratified the Qualitative test set by 9-mer overlap and report five retrained configurations in Table S3. All five scored lower in the no-overlap stratum, but their ordering was unchanged: BLOSUM62 (0.756) $<$ DeepNeo (0.861) $<$ Chai-1 (0.869) $<$ ESMC 300M (0.875) $<$ ESM3 Small (0.883). BLOSUM62 decreased by 0.115, roughly three times the 0.038–0.045 seen for Chai-1 and the two PLM configurations. Exact 15-mer matches were not the highest-scoring stratum (0.849–0.900 versus 0.871–0.932 for 9-mer overlap without an exact match). The two published percentile-rank tools reached 0.926 for NetMHCIIpan-4.3 and 0.914 for MixMHC2pred-2.0 in the no-overlap stratum; because they were not trained on our split, this stratum is not a train–test leakage control for them and these values are interpreted descriptively.

##### 1.5.4 Note on CD-HIT-based Retraining

We evaluated three 80% identity-filtered CD-HIT variants before adopting the stratified analysis. Even the best variant left 33% of test epitopes sharing a 9-mer with the filtered training set, compared with 81% before filtering, while reducing the training set from 112,871 to 35,348 entries (69%). Pooling positive and negative peptides during clustering also changed the negative-to-positive ratio from 0.72 to 0.20; class-specific clustering preserved the ratio but not the data loss.

Retraining on these splits would therefore change sequence redundancy and training-set size simultaneously, with class balance also affected in some variants. We used the stratified analysis above to quantify the residual overlap effect on the reported models.

#### 1.6 Serotype-Specific Performance Patterns

Per-serotype ROC-AUC values in the Qualitative dataset are given below for the five retrained configurations and the two percentile-rank tools. Serotypes differ at once in allele composition, assay composition, class balance, sequence diversity, and sample count, so this section reports the observed values and the ordering among them without assigning a cause to either.

##### 1.6.1 HLA-DP

HLA-DP gave the highest values of the four serotypes and the narrowest spread across representations: ESM3 Small 0.983, ESMC 300M 0.982, Chai-1 0.980, DeepNeo 0.962, and BLOSUM62 0.949, a range of 0.034. NetMHCIIpan-4.3 (0.963) falls inside that range, whereas MixMHC2pred-2.0 (0.947) lies just below it. HLA-DP also accounts for a larger share of the MS dataset (47.1%) than of the Qualitative dataset (16.9%).

##### 1.6.2 HLA-DQ

HLA-DQ separated the representations more widely than HLA-DP: ESMC 300M 0.893, ESM3 Small 0.887, and Chai-1 0.883, against 0.854 for DeepNeo and 0.839 for BLOSUM62, a range of 0.054. HLA-DQ contributes the smallest human share of the Qualitative dataset (11.9%).

##### 1.6.3 HLA-DR

HLA-DR contributes the largest share of the Qualitative dataset (69.3%) but was not the best-predicted serotype: ESM3 Small 0.905, ESMC 300M 0.901, Chai-1 0.895, DeepNeo 0.875, and BLOSUM62 0.847, a range of 0.058, the widest of the four serotypes. Sample count therefore does not reproduce the ordering across serotypes.

##### 1.6.4 Cross-Species Challenges

Murine H2 alleles gave the lowest per-serotype ROC-AUC values in the Qualitative dataset (0.813–0.871). In H2-out evaluation, pooled ROC-AUC ranged from 0.476 for BLOSUM62 to 0.682 for ESM3 Small, but H2-IAb accounts for 2,042 of the 3,120 H2 rows (65%). Equal weighting of the eight H2 molecules instead placed Chai-1 first (0.596), followed by ESMC 300M (0.586) and ESM3 Small (0.580). No pair survived Holm correction, and with only eight molecules this should be read as limited resolution rather than evidence of equivalence. NetMHCIIpan-4.3 and MixMHC2pred-2.0 score highly on H2 but both ship murine H-2 models, so H2-out is a species-out test only for the representations retrained here. These results describe transfer under the present protocol; identifying what information drives that transfer would require motif- or binding-core-level analysis.

#### 1.7 Data Imbalance and Under-Represented Alleles

Serotype-specific ROC-AUC does not increase monotonically with the amount of data available for each serotype. HLA-DR supplies 69.3% of the Qualitative dataset, HLA-DP 16.9% and HLA-DQ 11.9%, whereas the PLM configurations reach 0.982–0.983 on HLA-DP, 0.901–0.905 on HLA-DR and 0.887–0.893 on HLA-DQ. The serotype with the most data is therefore not the best predicted. We report this dissociation without assigning a cause: serotypes differ simultaneously in data volume, in the number of distinct alleles they contribute, in assay composition, and in positive/negative balance, and the present design does not separate these.

Under-represented alleles, particularly those from non-European populations and rare HLA variants, remain a challenge for pan-allelic prediction tools. The allele-wise analysis required at least five positives and five negatives, retaining 58 of 126 alleles while keeping 91.5% of test rows. LOMO withholds all data for one molecule and is the closest proxy available here for an allele absent from training. The analysis included 47 pair-named molecules with at least 20 positives, at least 20 negatives, and a minority-class fraction of at least 20% (38 $\beta$-chain units for DeepNeo). The proxy is therefore restricted to reasonably well-characterized molecules and does not directly measure performance on genuinely rare or sparsely characterized alleles.

#### 1.8 Detailed Software and Computational Environment

All structure-prediction models used in this study—AlphaFold 3, Chai-1, and Boltz-1—were executed locally based on publicly available source code. Since the original implementations did not support direct extraction of single and pair features during inference, we modified the code to expose the intermediate representations and bypass the diffusion modules. These modified versions were used exclusively for feature generation. The resulting scripts have been publicly released at `https://github.com/daylight-00/alphafold3`, `https://github.com/daylight-00/chai-lab`, and `https://github.com/daylight-00/boltz`, with all modifications traceable via commit history. Notably, AlphaFold 3 has since added official support for feature extraction.

Protein language model (PLM) embeddings were generated using ESM v3.1.6 (`https://doi.org/10.5281/zenodo.15020067`). For local inference, we used ESMC 300M and 600M models with FlashAttention v2.7.4.post1 (`https://github.com/Dao-AILab/flash-attention`). For larger models, including ESM3 Small/Medium/Large and ESMC 6B, we accessed the Forge API provided by EvolutionaryScale. All embeddings were generated on NVIDIA H100 SXM5 80GB GPUs, and stored in HDF5 format using h5py v3.13.0 (`https://github.com/h5py/h5py`).

Multiple sequence alignments (MSAs) were constructed for MHC sequences as required by structure prediction models. For AlphaFold 3 and Chai-1, MSAs were generated using jackhmmer from HMMER v3.4 (`http://hmmer.org/`) with the following sequence databases: BFD, UniRef90 (v2022_05), UniProt (v2021_04), and MGnify (v2022_05). For Boltz-1, MSAs were obtained via the ColabFold server (`https://api.colabfold.com`) using UniRef30 (v2023_02) and ColabFoldDB (v2021_08).

Model training and evaluation were performed in Python v3.13.2 (`https://www.python.org/`) using PyTorch v2.6.0 (`https://github.com/pytorch/pytorch`). All experiments were run on NVIDIA H100 SXM5 80GB GPUs. To ensure reproducibility, training was conducted under fixed random seeds and consistent hardware configurations.

For baseline comparisons, DeepNeo was re-implemented in PyTorch based on the official Theano-based source code and accompanying publications. Our implementation matched the original model architecture and training configuration, using stochastic gradient descent with momentum 0.9 and, following the learning-rate sweep described in the main text, a learning rate of $10^{-3}$. Benchmarking was performed under identical conditions to our proposed model. NetMHCIIpan-4.3 (`https://services.healthtech.dtu.dk/services/NetMHCIIpan-4.3/`) and MixMHC2pred-2.0, build 2.0.2 (`https://github.com/GfellerLab/MixMHC2pred`) were used in their official forms with default parameters, except that MixMHC2pred-2.0 was run with the `no_context` option to match our standardized input format. The NetMHCIIpan binary used for this study identifies itself as version **4.3g**. NetMHCIIpan-4.3 was scored with %Rank_EL for the Qualitative, MS, LOMO, H2-out, and serotype analyses and with %Rank_BA for the IC50 analyses. MixMHC2pred-2.0 has no affinity head and was scored on its single %Rank output. Percentile ranks are calibrated within alleles, so pooled and per-allele aggregation are interpreted separately. Both published tools were trained on IEDB or immunopeptidomics data that overlap the present test data. Their values are therefore used as contextual references rather than independent external benchmarks.

Dataset preprocessing, statistical analysis, and visualization were performed using NumPy v1.26.3 (`https://github.com/numpy/numpy`), pandas v2.2.3 (`https://github.com/pandas-dev/`

pandas), scikit-learn v1.6.1 (https://github.com/scikit-learn/scikit-learn), Matplotlib v3.10.1 (https://github.com/matplotlib/matplotlib), and seaborn v0.13.2 (https://github.com/mwaskom/seaborn). Some tabular processing and manual review were conducted in Microsoft Excel (Microsoft 365).

Human MHC allele sequences were retrieved from the IPD-IMGT/HLA database, release 3.59.0 (https://github.com/ANHIG/IMGTHLA) and murine H2 sequences from UniProt (https://www.uniprot.org/), as described in Section 1.1. Epitope–MHC binding records were obtained from the IEDB Database Export v3 (dated 2025/04/21; https://www.iedb.org/database_export_v3.php). For MSA construction, we used the following databases: BFD, UniRef90, UniProt, and MGnify, in accordance with each model's requirements.

##### 1.9 Reference Tool Allele Coverage

Neither published tool covers every allele in our benchmark, so each was scored on the test rows carrying an allele it actually supports.

NetMHCIIpan-4.3's pseudosequence table contains eight murine H-2 alleles but not H2-IAg7, which excludes 176 cases in the Qualitative test set, 158 in MS and 40 in IC50. MixMHC2pred-2.0 ships position-weight matrices for thirteen murine H-2 alleles alongside its human set, including H2-IAg7, and so covers every allele present in the Qualitative, MS and IC50 test sets. One further Qualitative test row is unscored by both tools for a reason unrelated to allele coverage: the peptide XPLALQFAELPVNKG (H2-IAk) contains a non-standard residue. NetMHCIIpan-4.3's Qualitative sample size is therefore 48,352 less the 176 H2-IAg7 rows and this one row.

The resulting effective sample sizes are, for NetMHCIIpan-4.3 and MixMHC2pred-2.0 respectively: Qualitative 48,175 and 48,351 of 48,352; MS 33,332 and 33,489 of 33,490; IC50 14,110 and 14,150 of 14,150. Every PREpiBind configuration and DeepNeo is scored on the full test set. Supplementary Table S14 repeats these alongside the values.

##### 1.10 Use of Generative AI Tools

Claude Opus 5 was used for language editing and refinement of author-prepared drafts, assistance with validation of analysis code, and consistency checks among the manuscript, supplementary materials, and underlying data. All AI-assisted outputs were reviewed and verified by the authors, who take full responsibility for the final content, analyses, and conclusions.

### 2 Supplementary Tables

#### 2.1 Sample counts and sequence overlap

**Table S1:** Number of samples in train and test sets for each dataset.

| Dataset | Train | Test | Total |
|---|---|---|---|
| Qualitative | 112,871 | 48,352 | 161,223 |
| IC50 | 33,004 | 14,150 | 47,154 |
| MS | 77,954 | 33,490 | 111,444 |

**Table S2:** Overlap between unique test and training epitopes in the Qualitative dataset (26,737 unique test epitopes; 46,487 unique training epitopes).

| **Overlap category** | **Count** | **%** |
|---|---|---|
| No 9-mer overlap with training | 4,967 | 18.6% |
| ≥1 9-mer shared with training | 21,770 | 81.4% |
| of which: exact 15-mer match | 15,628 | 58.5% |
| of which: 9-mer overlap, not exact match | 6,142 | 23.0% |
| 15-mer identity ≥80% (≤3 mismatches) | 17,520 | 65.5% |
| 15-mer identity ≥86.7% (≤2 mismatches) | 17,120 | 64.0% |
| 15-mer identity ≥93.3% (≤1 mismatch) | 16,722 | 62.5% |

**Table S3:** Stratified ROC-AUC on the Qualitative test set by 9-mer overlap with training. Values are means over three seeds. Strata are defined over test rows; DeepNeo uses the $\beta$-chain split and therefore has slightly smaller counts.

| **Stratum** | **n** | **BLOSUM62** | **DeepNeo†** | **Chai-1** | **ESMC 300M** | **ESM3 Small** |
|---|---|---|---|---|---|---|
| Overall test set | 47,666–48,352 | 0.871 | 0.894 | 0.914 | 0.919 | 0.921 |
| No 9-mer overlap | 5,222–5,228 | 0.756 | 0.861 | 0.869 | 0.875 | 0.883 |
| ≥1 9-mer overlap | 42,444–43,124 | 0.864 | 0.882 | 0.905 | 0.911 | 0.912 |
| Exact 15-mer match | 35,473–36,146 | 0.849 | 0.867 | 0.894 | 0.899 | 0.900 |
| 9-mer overlap, not exact | 6,971–6,978 | 0.871 | 0.906 | 0.917 | 0.926 | 0.932 |

#### 2.2 Domain boundary coordinates and UniProt IDs

**Table S4:** Domain boundary coordinates and UniProt IDs for each MHC class II gene.

| **Gene** | **Domain Position** | **UniProt ID** |
|---|---|---|
| HLA-DPA1 | 35–114 | P20036 |
| HLA-DPB1 | 42–114 | P04440 |
| HLA-DQA1 | 29–108 | P01909 |
| HLA-DQB1 | 45–119 | P01920 |
| HLA-DRA | 30–109 | P01903 |
| HLA-DRB1 | 42–116 | P01911 |
| H2-IAb ($\alpha$) | 30–110 | P14434 |
| H2-IAb ($\beta$) | 40–114 | P14483 |
| H2-IAd ($\alpha$) | 30–110 | P04228 |
| H2-IAd ($\beta$) | 40–114 | P01921 |
| H2-IAg7 ($\alpha$) | 30–110 | P04228 |
| H2-IAg7 ($\beta$) | 40–112 | Q31135 |
| H2-IAk ($\alpha$) | 30–110 | P01910 |
| H2-IAk ($\beta$) | 40–112 | P06343 |
| H2-IAq ($\alpha$) | 1–76 | P04227 |
| H2-IAq ($\beta$) | 40–114 | P06342 |
| H2-IAs ($\alpha$) | 7–87 | P14437 |
| H2-IAs ($\beta$) | 40–112 | P06345 |
| H2-IAu ($\alpha$) | 1–81 | P14438 |
| H2-IAu ($\beta$) | 40–112 | P06344 |
| H2-IEd ($\alpha$) | 30–109 | P01904 |
| H2-IEd ($\beta$) | 40–114 | P01915 |
| H2-IEk ($\alpha$) | 30–109 | P04224 |
| H2-IEk ($\beta$) | 39–113 | Q31163 |

#### 2.3 Embedding dimensions

**Table S5:** Output dimensions of each protein representation method. $L$ denotes the input sequence length.

| Method | Original Features | Final Embedding |
|---|---|---|
| BLOSUM62 | $L \times 25$ | $L \times 25$ |
| AlphaFold 3 | $L \times 384$, $L \times L \times 128$ | $L \times 640$ |
| Boltz-1 | $L \times 384$, $L \times L \times 128$ | $L \times 640$ |
| Chai-1 | $L \times 384$, $L \times L \times 256$ | $L \times 896$ |
| ESMC 300M | $L \times 960$ | $L \times 960$ |
| ESMC 600M | $L \times 1152$ | $L \times 1152$ |
| ESMC 6B | $L \times 2560$ | $L \times 2560$ |
| ESM3 Small | $L \times 1536$ | $L \times 1536$ |
| ESM3 Medium | $L \times 2560$ | $L \times 2560$ |
| ESM3 Large | $L \times 6144$ | $L \times 6144$ |

#### 2.4 Model parameters

**Table S6:** Number of trainable model parameters for each embedding method.

| Embedding Model | Model Parameters |
|---|---|
| BLOSUM62 | 39,450 |
| AlphaFold 3 | 24,823,041 |
| Boltz-1 | 24,823,041 |
| Chai-1 | 48,629,505 |
| ESMC 300M | 55,820,161 |
| ESMC 600M | 80,365,825 |
| ESMC 6B | 396,661,761 |
| ESM3 Small | 142,838,785 |
| ESM3 Medium | 396,661,761 |
| ESM3 Large | 2,284,204,033 |
| DeepNeo (baseline) | 585,311 |

#### 2.5 Full benchmark performance metrics

Performance metrics for all models and datasets. Values for trained models are reported as mean $\pm$ standard deviation across three seeds; aggregation and decision thresholds are described in the Methods. For percentile-rank tools, only ROC-AUC and PR-AUC are reported (—). Bold values indicate the best result per column within each model class. Every representation is evaluated on every dataset; Figure 2(b)–(d) of the main text plots a subset of them.

**Table S7:** Benchmark performance on the Qualitative dataset.

| Method | ROC-AUC | PR-AUC | F1 | Accuracy | MCC |
|---|---|---|---|---|---|
| BLOSUM62 | 0.871 ±0.005 | 0.907 ±0.004 | 0.818 ±0.003 | 0.788 ±0.004 | 0.565 ±0.009 |
| AlphaFold 3 | 0.876 ±0.006 | 0.913 ±0.004 | 0.823 ±0.005 | 0.791 ±0.006 | 0.569 ±0.013 |
| Boltz-1 | 0.884 ±0.004 | 0.918 ±0.003 | 0.830 ±0.002 | 0.799 ±0.003 | 0.587 ±0.008 |
| Chai-1 | 0.914 ±0.001 | 0.941 ±0.001 | 0.855 ±0.002 | 0.831 ±0.001 | 0.654 ±0.002 |
| ESMC 300M | 0.919 ±0.002 | 0.945 ±0.001 | 0.859 ±0.002 | 0.838 ±0.003 | 0.668 ±0.007 |
| ESMC 600M | 0.921 ±0.001 | 0.946 ±0.001 | 0.860 ±0.002 | 0.839 ±0.002 | 0.672 ±0.005 |
| ESMC 6B | 0.909 ±0.001 | 0.938 ±0.001 | 0.847 ±0.003 | 0.826 ±0.002 | 0.646 ±0.001 |
| ESM3 Small | 0.921 ±0.001 | 0.946 ±0.001 | 0.860 ±0.003 | 0.839 ±0.002 | 0.672 ±0.002 |
| ESM3 Medium | 0.920 ±0.003 | 0.945 ±0.002 | 0.859 ±0.004 | 0.837 ±0.004 | 0.668 ±0.008 |
| ESM3 Large | **0.927 ±0.002** | **0.951 ±0.001** | **0.867 ±0.001** | **0.847 ±0.002** | **0.686 ±0.006** |
| DeepNeo† | 0.894 ±0.002 | 0.927 ±0.001 | 0.838 ±0.002 | 0.811 ±0.002 | 0.612 ±0.004 |
| NetMHCIIpan-4.3† | 0.838 | 0.898 | — | — | — |
| MixMHC2pred-2.0† | 0.815 | 0.882 | — | — | — |

**Table S8:** Benchmark performance on the MS dataset. Percentile-rank tools are scored on the test rows carrying supported alleles; effective sample sizes are reported in Table S14.

| Method | ROC-AUC | PR-AUC | F1 | Accuracy | MCC |
|---|---|---|---|---|---|
| BLOSUM62 | 0.959 ±0.007 | 0.945 ±0.009 | 0.864 ±0.009 | 0.895 ±0.008 | 0.780 ±0.017 |
| AlphaFold 3 | 0.949 ±0.004 | 0.932 ±0.005 | 0.847 ±0.008 | 0.881 ±0.006 | 0.749 ±0.013 |
| Boltz-1 | 0.962 ±0.001 | 0.950 ±0.002 | 0.871 ±0.004 | 0.897 ±0.005 | 0.787 ±0.009 |
| Chai-1 | 0.987 ±0.002 | 0.982 ±0.002 | 0.934 ±0.005 | 0.948 ±0.004 | 0.892 ±0.009 |
| ESMC 300M | 0.990 ±0.001 | 0.987 ±0.001 | 0.944 ±0.003 | 0.956 ±0.002 | 0.908 ±0.005 |
| ESMC 600M | 0.990 ±0.000 | 0.987 ±0.001 | 0.944 ±0.001 | 0.956 ±0.001 | 0.908 ±0.002 |
| ESMC 6B | 0.987 ±0.001 | 0.982 ±0.001 | 0.931 ±0.002 | 0.946 ±0.002 | 0.888 ±0.004 |
| ESM3 Small | 0.990 ±0.000 | 0.987 ±0.001 | 0.947 ±0.002 | 0.958 ±0.002 | 0.913 ±0.004 |
| ESM3 Medium | 0.990 ±0.000 | 0.987 ±0.000 | 0.947 ±0.002 | 0.959 ±0.002 | 0.914 ±0.003 |
| ESM3 Large | **0.994 ±0.001** | **0.992 ±0.001** | **0.957 ±0.003** | **0.966 ±0.002** | **0.930 ±0.004** |
| DeepNeo† | 0.980 ±0.000 | 0.974 ±0.001 | 0.917 ±0.000 | 0.935 ±0.000 | 0.864 ±0.001 |
| NetMHCIIpan-4.3† | 0.974 | 0.971 | — | — | — |
| MixMHC2pred-2.0† | 0.951 | 0.945 | — | — | — |

**Table S9:** Benchmark performance on the IC50 dataset (<500 nM threshold).

| Method | ROC-AUC | PR-AUC | F1 | Accuracy | MCC |
|---|---|---|---|---|---|
| BLOSUM62 | 0.801 ±0.003 | 0.687 ±0.003 | 0.622 ±0.012 | 0.739 ±0.003 | 0.427 ±0.010 |
| AlphaFold 3 | 0.803 ±0.003 | 0.694 ±0.004 | 0.634 ±0.001 | 0.742 ±0.003 | 0.436 ±0.005 |
| Boltz-1 | 0.807 ±0.005 | 0.700 ±0.005 | 0.647 ±0.003 | 0.747 ±0.004 | 0.451 ±0.005 |
| Chai-1 | 0.827 ±0.005 | 0.730 ±0.007 | 0.662 ±0.006 | 0.762 ±0.003 | 0.481 ±0.007 |
| ESMC 300M | 0.838 ±0.001 | 0.743 ±0.002 | 0.676 ±0.005 | 0.769 ±0.002 | 0.498 ±0.005 |
| ESMC 600M | 0.837 ±0.001 | 0.743 ±0.001 | 0.675 ±0.003 | 0.770 ±0.000 | 0.499 ±0.002 |
| ESMC 6B | 0.825 ±0.001 | 0.729 ±0.001 | 0.658 ±0.004 | 0.762 ±0.001 | 0.479 ±0.003 |
| ESM3 Small | **0.839 ±0.002** | **0.744 ±0.002** | **0.682 ±0.009** | **0.771 ±0.002** | **0.504 ±0.007** |
| ESM3 Medium | 0.835 ±0.002 | 0.742 ±0.001 | 0.675 ±0.005 | 0.768 ±0.001 | 0.496 ±0.003 |
| ESM3 Large | 0.835 ±0.001 | 0.742 ±0.002 | 0.667 ±0.005 | 0.768 ±0.002 | 0.494 ±0.005 |
| DeepNeo† | 0.830 ±0.001 | 0.732 ±0.002 | 0.664 ±0.002 | 0.763 ±0.001 | 0.482 ±0.001 |
| NetMHCIIpan-4.3† | 0.909 | 0.851 | — | — | — |
| MixMHC2pred-2.0† | 0.714 | 0.590 | — | — | — |

**Table S10:** Benchmark performance on the IC50 dataset (<1,000 nM threshold).

| Method | ROC-AUC | PR-AUC | F1 | Accuracy | MCC |
|---|---|---|---|---|---|
| BLOSUM62 | 0.796 ±0.001 | 0.754 ±0.001 | 0.695 ±0.002 | 0.722 ±0.001 | 0.440 ±0.000 |
| AlphaFold 3 | 0.796 ±0.009 | 0.757 ±0.010 | 0.695 ±0.010 | 0.722 ±0.007 | 0.439 ±0.015 |
| Boltz-1 | 0.806 ±0.001 | 0.772 ±0.003 | 0.707 ±0.002 | 0.731 ±0.001 | 0.460 ±0.001 |
| Chai-1 | 0.823 ±0.001 | 0.790 ±0.001 | 0.722 ±0.004 | 0.744 ±0.001 | 0.485 ±0.002 |
| ESMC 300M | 0.835 ±0.002 | 0.802 ±0.004 | **0.740 ±0.002** | 0.755 ±0.001 | **0.510 ±0.003** |
| ESMC 600M | 0.835 ±0.002 | 0.806 ±0.002 | 0.736 ±0.006 | 0.755 ±0.001 | 0.509 ±0.003 |
| ESMC 6B | 0.825 ±0.003 | 0.795 ±0.003 | 0.719 ±0.006 | 0.746 ±0.003 | 0.488 ±0.006 |
| ESM3 Small | 0.834 ±0.004 | 0.802 ±0.005 | 0.735 ±0.004 | 0.754 ±0.003 | 0.506 ±0.007 |
| ESM3 Medium | 0.832 ±0.002 | 0.801 ±0.002 | 0.730 ±0.004 | 0.754 ±0.001 | 0.504 ±0.001 |
| ESM3 Large | **0.836 ±0.002** | **0.806 ±0.002** | 0.729 ±0.004 | **0.756 ±0.002** | 0.508 ±0.004 |
| DeepNeo† | 0.822 ±0.002 | 0.790 ±0.003 | 0.722 ±0.003 | 0.746 ±0.002 | 0.488 ±0.005 |
| NetMHCIIpan-4.3† | 0.899 | 0.880 | — | — | — |
| MixMHC2pred-2.0† | 0.702 | 0.664 | — | — | — |

#### 2.6 Bootstrap confidence intervals

**Table S11:** Bootstrap uncertainty on pooled Qualitative ROC-AUC. The 95% intervals were estimated from 1,000 test-set resamples. Pairwise differences use the same resampled rows for both configurations. Seed-level standard deviations are reported separately and reflect training variability.

| **Configuration** | **ROC-AUC** | **95% CI** |
|---|---|---|
| BLOSUM62 | 0.871 | [0.869, 0.874] |
| DeepNeo† | 0.894 | [0.892, 0.897] |
| Chai-1 | 0.914 | [0.912, 0.916] |
| ESMC 300M | 0.919 | [0.917, 0.921] |
| ESM3 Small | 0.921 | [0.919, 0.923] |
| *Paired differences* | | |
| **Comparison** | **Δ** | **95% CI** |
| BLOSUM62 − ESM3 Small | -0.0498 | [-0.0516, -0.0478] |
| BLOSUM62 − ESMC 300M | -0.0479 | [-0.0497, -0.0459] |
| BLOSUM62 − Chai-1 | -0.0425 | [-0.0444, -0.0405] |
| Chai-1 − ESM3 Small | -0.0073 | [-0.0083, -0.0061] |
| Chai-1 − ESMC 300M | -0.0055 | [-0.0065, -0.0044] |
| ESM3 Small − ESMC 300M | +0.0018 | [+0.0009, +0.0028] |

#### 2.7 UMAP cluster separation

**Table S12:** Silhouette coefficients (cosine distance) for MHC-II gene-family labels computed from per-allele mean-pooled embeddings. Macro, micro, and median summaries are reported for all ten representations. Asterisks mark the four representative methods displayed in Supplementary Figure S2.

| **Method** | **Silhouette (macro)** | **Micro** | **Median** |
|---|---|---|---|
| BLOSUM62* | 0.55 | 0.39 | 0.40 |
| AlphaFold 3 | 0.05 | 0.02 | −0.04 |
| Boltz-1 | 0.49 | 0.39 | 0.40 |
| Chai-1* | 0.56 | 0.51 | 0.58 |
| ESMC 300M* | 0.51 | 0.57 | 0.69 |
| ESMC 600M | 0.67 | 0.68 | 0.75 |
| ESMC 6B | 0.71 | 0.63 | 0.66 |
| ESM3 Small* | 0.53 | 0.33 | 0.19 |
| ESM3 Medium | 0.60 | 0.62 | 0.71 |
| ESM3 Large | 0.66 | 0.58 | 0.62 |

#### 2.8 Per-allele versus pooled ROC-AUC

**Table S13:** Per-allele versus pooled ROC-AUC on the Qualitative dataset. Per-allele values use every allele supported by each method; the main text and Figure 3a use the shared paired convention in Table S15. Pooled ROC-AUC is higher for every representation.

| **Method** | **Per-allele** | **Pooled** |
|---|---|---|
| BLOSUM62 | 0.780 | 0.871 |
| Chai-1 | 0.843 | 0.914 |
| ESMC 300M | 0.852 | 0.919 |
| ESM3 Small | 0.851 | 0.921 |
| DeepNeo† | 0.811 | 0.894 |
| NetMHCIIpan-4.3† | 0.795 | 0.838 |
| MixMHC2pred-2.0† | 0.774 | 0.815 |

**Table S14:** Pooled ROC-AUC and effective sample sizes for all representations and the two published tools. The tools are scored only on supported alleles and were not retrained on our splits, so their values are contextual rather than like-for-like. NetMHCIIpan-4.3 uses its EL rank outside IC50 and BA rank for IC50; MixMHC2pred-2.0 uses its percentile-rank output.

| | **Qualitative** | **MS** | **IC50 <500** | **IC50 <1000** | **H2-Out** |
|---|---|---|---|---|---|
| *Pooled ROC-AUC* | | | | | |
| BLOSUM62 | 0.871 | 0.959 | 0.801 | 0.796 | 0.476 |
| Chai-1 | 0.914 | 0.987 | 0.827 | 0.823 | 0.628 |
| ESMC 300M | 0.919 | 0.990 | 0.838 | 0.835 | 0.680 |
| ESM3 Small | 0.921 | 0.990 | 0.839 | 0.834 | 0.682 |
| DeepNeo | 0.894 | 0.980 | 0.830 | 0.822 | 0.529 |
| *NetMHCIIpan-4.3* | 0.838 | 0.974 | 0.909 | 0.899 | 0.883 |
| *MixMHC2pred-2.0* | 0.815 | 0.951 | 0.714 | 0.702 | 0.875 |
| *Effective n. The retrained models are scored on every test row.* | | | | | |
| *NetMHCIIpan-4.3* | 48,175 | 33,332 | 14,110 | 14,110 | 2,557 |
| *MixMHC2pred-2.0* | 48,351 | 33,489 | 14,150 | 14,150 | 3,119 |

**Table S15:** Qualitative per-allele ROC-AUC under three unit conventions. **all** uses every allele supported by each method; **paired** uses the 57 $\alpha/\beta$ alleles shared by the pair-named methods and is used in Figure 3a; **beta** collapses to the $\beta$ chain so DeepNeo can be paired with the others. Differences between conventions are small and do not support an ordering between ESMC 300M and ESM3 Small.

| | **all** | $n$ | **paired** | **beta** | $n$ **(beta)** | **all − beta** |
|---|---|---|---|---|---|---|
| BLOSUM62 | 0.780 | 58 | 0.780 | 0.778 | 48 | +0.002 |
| Chai-1 | 0.843 | 58 | 0.843 | 0.843 | 48 | +0.000 |
| ESMC 300M | 0.852 | 58 | 0.852 | 0.851 | 48 | +0.001 |
| ESM3 Small | 0.851 | 58 | 0.852 | 0.851 | 48 | +0.000 |
| DeepNeo | 0.811 | 48 | 0.811 | 0.811 | 48 | +0.000 |
| *NetMHCIIpan-4.3* | 0.795 | 57 | 0.795 | 0.801 | 47 | -0.006 |
| *MixMHC2pred-2.0* | 0.774 | 58 | 0.772 | 0.783 | 48 | -0.009 |

**Table S16:** Mean per-molecule ROC-AUC. The Per Molecule column uses the allele unit of Figure 3a; LOMO and H2-out use the $\beta$-chain unit. Published tools were not retrained under the leave-out splits and are shown only as contextual references. NetMHCIIpan-4.3 lacks H2-IAg7 coverage and therefore contributes seven H2-out molecules. Counts are given in Table S17.

| | **Per Molecule** | **LOMO – Whole** | **LOMO – H2** | **H2-Out** |
|---|---|---|---|---|
| BLOSUM62 | 0.780 | 0.713 | 0.573 | 0.470 |
| Chai-1 | 0.843 | 0.792 | 0.660 | 0.596 |
| ESMC 300M | 0.852 | 0.796 | 0.666 | 0.586 |
| ESM3 Small | 0.852 | 0.778 | 0.618 | 0.580 |
| DeepNeo | 0.811 | 0.761 | 0.560 | 0.491 |
| *NetMHCIIpan-4.3* | 0.795 | 0.805 | 0.780 | 0.670 |
| *MixMHC2pred-2.0* | 0.772 | 0.779 | 0.759 | 0.740 |

**Table S17:** Number of molecules behind each mean in Table S16.

| | **Per Molecule** | **LOMO – Whole** | **LOMO – H2** | **H2-Out** |
|---|---|---|---|---|
| BLOSUM62 | 57 | 38 | 3 | 8 |
| Chai-1 | 57 | 38 | 3 | 8 |
| ESMC 300M | 57 | 38 | 3 | 8 |
| ESM3 Small | 57 | 38 | 3 | 8 |
| DeepNeo | 48 | 38 | 3 | 8 |
| *NetMHCIIpan-4.3* | 57 | 38 | 3 | 7 |
| *MixMHC2pred-2.0* | 57 | 38 | 3 | 8 |

**Table S18:** Holm-corrected paired Wilcoxon tests for the 57 shared allele pairs in Figure 3a. $\Delta$ is the median paired difference $(a - b)$. DeepNeo is excluded at this unit because it models only the $\beta$ chain; its comparisons are reported in Table S19. Published-tool comparisons are included for completeness but should be interpreted in light of their overlapping training data.

| | $n$ | $\Delta$ | $p$ | $p_{\text{adj}}$ | **Sig.** |
|---|---|---|---|---|---|
| BLOSUM62 vs Chai-1 | 57 | -0.063 | 1.02e-09 | 1.46e-08 | *** |
| BLOSUM62 vs ESMC 300M | 57 | -0.069 | 9.73e-10 | 1.46e-08 | *** |
| BLOSUM62 vs ESM3 Small | 57 | -0.071 | 9.73e-10 | 1.46e-08 | *** |
| BLOSUM62 vs NetMHCIIpan-4.3 | 57 | -0.002 | 3.76e-01 | 9.11e-01 | ns |
| BLOSUM62 vs MixMHC2pred-2.0 | 57 | +0.015 | 3.04e-01 | 9.11e-01 | ns |
| Chai-1 vs ESMC 300M | 57 | -0.008 | 2.32e-06 | 1.86e-05 | *** |
| Chai-1 vs ESM3 Small | 57 | -0.009 | 1.75e-05 | 9.10e-05 | *** |
| Chai-1 vs NetMHCIIpan-4.3 | 57 | +0.053 | 6.54e-05 | 2.62e-04 | *** |
| Chai-1 vs MixMHC2pred-2.0 | 57 | +0.077 | 9.67e-07 | 8.70e-06 | *** |
| ESMC 300M vs ESM3 Small | 57 | -0.000 | 7.18e-01 | 9.11e-01 | ns |
| ESMC 300M vs NetMHCIIpan-4.3 | 57 | +0.056 | 1.52e-05 | 9.10e-05 | *** |
| ESMC 300M vs MixMHC2pred-2.0 | 57 | +0.089 | 2.57e-07 | 2.83e-06 | *** |
| ESM3 Small vs NetMHCIIpan-4.3 | 57 | +0.062 | 9.80e-06 | 6.86e-05 | *** |
| ESM3 Small vs MixMHC2pred-2.0 | 57 | +0.087 | 1.24e-07 | 1.49e-06 | *** |
| NetMHCIIpan-4.3 vs MixMHC2pred-2.0 | 57 | +0.019 | 7.58e-07 | 7.58e-06 | *** |

**Table S19:** The same paired comparison after collapsing all methods onto the $\beta$-chain unit ($n = 47$), which permits direct comparison with DeepNeo. $\Delta$ is the median paired difference $(a-b)$. Published-tool comparisons remain contextual because those tools were not trained on the present splits.

| | $n$ | $\Delta$ | $p$ | $p_{\text{adj}}$ | **Sig.** |
|---|---|---|---|---|---|
| BLOSUM62 vs Chai-1 | 47 | -0.061 | 3.50e-10 | 6.66e-09 | *** |
| BLOSUM62 vs ESMC 300M | 47 | -0.068 | 3.09e-10 | 6.49e-09 | *** |
| BLOSUM62 vs ESM3 Small | 47 | -0.071 | 3.09e-10 | 6.49e-09 | *** |
| BLOSUM62 vs DeepNeo | 47 | -0.040 | 1.85e-06 | 2.04e-05 | *** |
| BLOSUM62 vs NetMHCIIpan-4.3 | 47 | -0.017 | 1.44e-01 | 4.32e-01 | ns |
| BLOSUM62 vs MixMHC2pred-2.0 | 47 | +0.006 | 7.57e-01 | 7.57e-01 | ns |
| Chai-1 vs ESMC 300M | 47 | -0.006 | 3.80e-07 | 5.70e-06 | *** |
| Chai-1 vs ESM3 Small | 47 | -0.010 | 2.12e-06 | 2.04e-05 | *** |
| Chai-1 vs DeepNeo | 47 | +0.023 | 1.85e-06 | 2.04e-05 | *** |
| Chai-1 vs NetMHCIIpan-4.3 | 47 | +0.050 | 1.68e-04 | 1.01e-03 | ** |
| Chai-1 vs MixMHC2pred-2.0 | 47 | +0.063 | 1.61e-06 | 1.94e-05 | *** |
| ESMC 300M vs ESM3 Small | 47 | -0.004 | 2.95e-01 | 5.91e-01 | ns |
| ESMC 300M vs DeepNeo | 47 | +0.030 | 4.98e-08 | 8.46e-07 | *** |
| ESMC 300M vs NetMHCIIpan-4.3 | 47 | +0.050 | 2.52e-05 | 1.76e-04 | *** |
| ESMC 300M vs MixMHC2pred-2.0 | 47 | +0.075 | 5.13e-07 | 7.19e-06 | *** |
| ESM3 Small vs DeepNeo | 47 | +0.034 | 9.46e-09 | 1.70e-07 | *** |
| ESM3 Small vs NetMHCIIpan-4.3 | 47 | +0.060 | 1.79e-05 | 1.43e-04 | *** |
| ESM3 Small vs MixMHC2pred-2.0 | 47 | +0.077 | 2.04e-07 | 3.27e-06 | *** |
| DeepNeo vs NetMHCIIpan-4.3 | 47 | +0.032 | 4.42e-02 | 1.77e-01 | ns |
| DeepNeo vs MixMHC2pred-2.0 | 47 | +0.052 | 3.57e-04 | 1.79e-03 | ** |
| NetMHCIIpan-4.3 vs MixMHC2pred-2.0 | 47 | +0.019 | 5.53e-07 | 7.19e-06 | *** |

**Table S20:** Holm-corrected paired Wilcoxon tests for the 38 withheld molecules in Figure 3b (LOMO), at the $\beta$-chain unit so DeepNeo is comparable with the others. $\Delta$ is the median paired difference $(a-b)$. The published tools are absent because they were not retrained under the leave-one-molecule-out splits.

| | $n$ | $\Delta$ | $p$ | $p_{\text{adj}}$ | **Sig.** |
|---|---|---|---|---|---|
| BLOSUM62 vs Chai-1 | 38 | -0.077 | 7.28e-12 | 7.28e-11 | *** |
| BLOSUM62 vs ESMC 300M | 38 | -0.085 | 7.28e-12 | 7.28e-11 | *** |
| BLOSUM62 vs ESM3 Small | 38 | -0.062 | 1.38e-10 | 1.11e-09 | *** |
| BLOSUM62 vs DeepNeo | 38 | -0.049 | 1.60e-05 | 1.12e-04 | *** |
| Chai-1 vs ESMC 300M | 38 | -0.003 | 8.71e-02 | 1.30e-01 | ns |
| Chai-1 vs ESM3 Small | 38 | +0.012 | 4.47e-03 | 1.34e-02 | * |
| Chai-1 vs DeepNeo | 38 | +0.022 | 1.74e-04 | 8.70e-04 | *** |
| ESMC 300M vs ESM3 Small | 38 | +0.014 | 2.14e-04 | 8.70e-04 | *** |
| ESMC 300M vs DeepNeo | 38 | +0.024 | 3.95e-05 | 2.37e-04 | *** |
| ESM3 Small vs DeepNeo | 38 | +0.006 | 6.51e-02 | 1.30e-01 | ns |

### 3 Supplementary Figures

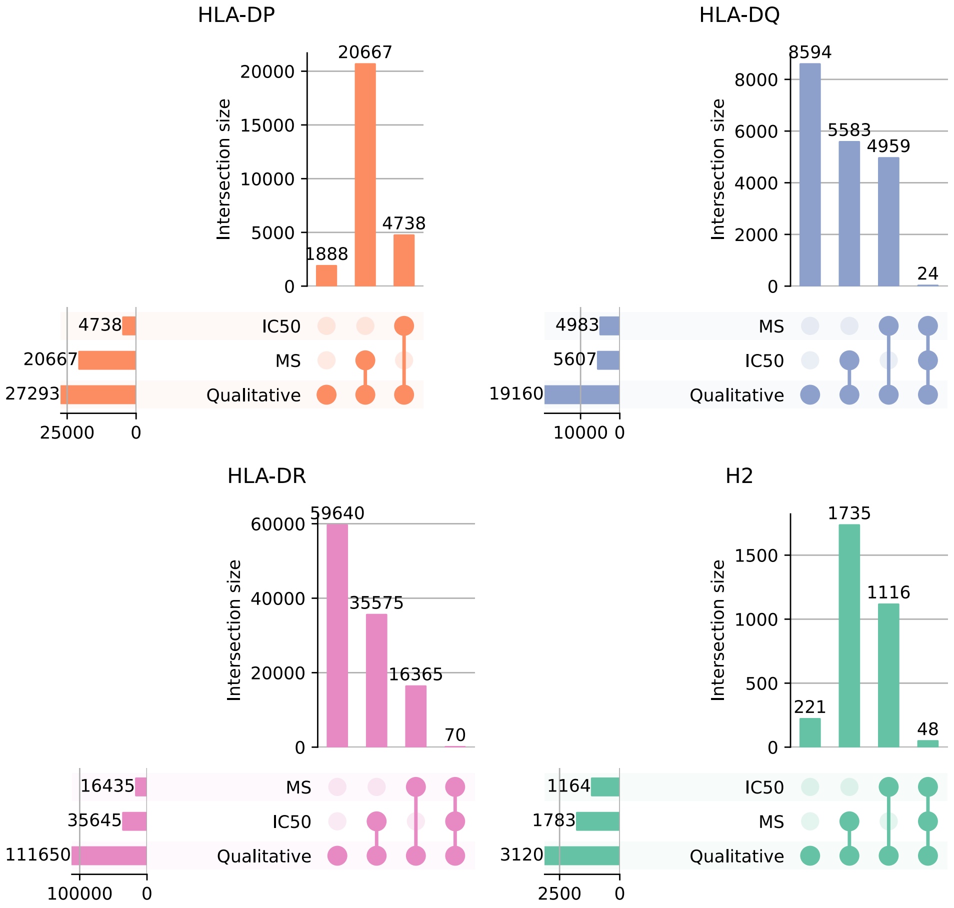


**Figure S1:** Upset plot overlaps among different MHC serotypes (HLA-DR, HLA-DQ, HLA-DP, and H2) for each dataset type. Intersection sizes are indicated by bar heights, and dataset types involved in each intersection are marked by connected dots below each bar. Intersections are shown at the MHC–epitope level.

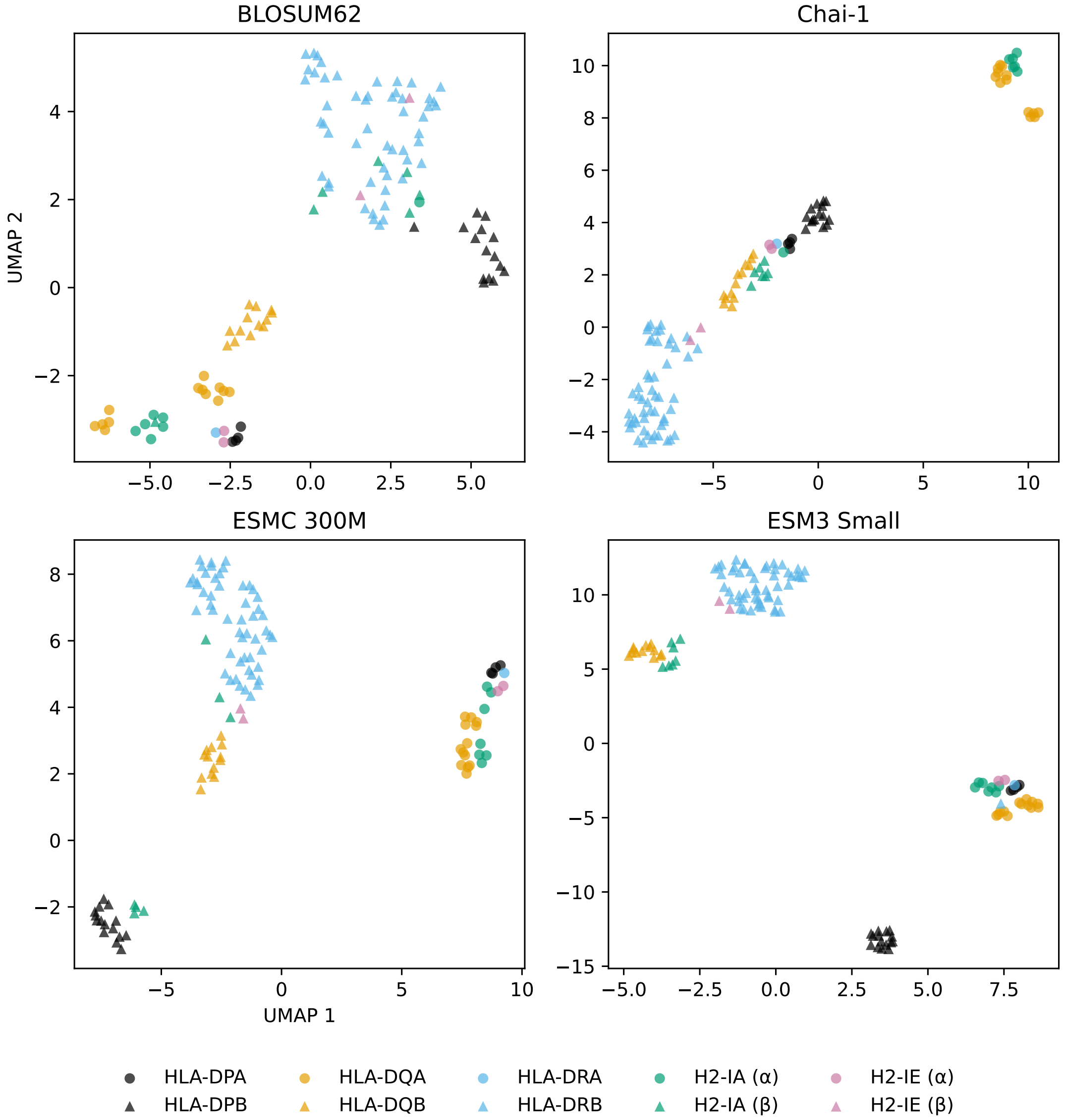


**Figure S2: UMAP visualization of MHC-II embeddings across representation methods.** Two-dimensional UMAP projections of MHC $\alpha$- and $\beta$-chain embeddings are shown for BLOSUM62, Chai-1, ESMC 300M, and ESM3 Small. Each point represents a unique MHC allele, colored by gene family: HLA-DPA (blue), HLA-DQA (green), HLA-DRA (orange), HLA-DPB (purple), HLA-DQB (red), HLA-DRB (brown), H2-IA (pink), and H2-IE (cyan). Macro-averaged silhouette coefficients (cosine distance) were 0.55 for BLOSUM62, 0.56 for Chai-1, 0.51 for ESMC 300M, and 0.53 for ESM3 Small.

#### References

[1] Jonas B Nilsson, Saghar Kaabinejadian, Hooman Yari, Michel GD Kester, Peter van Balen, William H Hildebrand, and Morten Nielsen. Accurate prediction of HLA class II antigen presentation across all loci using tailored data acquisition and refined machine learning. *Science Advances*, 9(47):eadj6367, 2023.

[2] Julien Racle, Philippe Guillaume, Julien Schmidt, Justine Michaux, Amédé Larabi, Kelvin Lau, Marta AS Perez, Giancarlo Croce, Raphaël Genolet, George Coukos, et al. Machine learning predictions of MHC-II specificities reveal alternative binding mode of class II epitopes. *Immunity*, 56(6):1359–1375, 2023.

[3] Leland McInnes, John Healy, and James Melville. UMAP: Uniform manifold approximation and projection. *Journal of Open Source Software*, 3(29):861, 2018. doi: 10.21105/joss.00861.

[4] Yohan Kim, John Sidney, Clemencia Pinilla, Alessandro Sette, and Bjoern Peters. Dataset size and composition impact the reliability of performance benchmarks for peptide–MHC binding predictions. *BMC Bioinformatics*, 15:241, 2014. doi: 10.1186/1471-2105-15-241.

[5] Birkir Reynisson, Bruno Alvarez, Sinu Paul, Bjoern Peters, and Morten Nielsen. NetMHCpan-4.1 and NetMHCIIpan-4.0: improved predictions of MHC antigen presentation by concurrent motif deconvolution and integration of ms MHC eluted ligand data. *Nucleic Acids Research*, 48(W1):W449–W454, 2020.
